# *In-Silico* Analysis of Secondary Metabolites that Modulates Enzymes of Cholesterol Target

**DOI:** 10.1101/2020.07.24.219998

**Authors:** Rishab Marahatha, Saroj Basnet, Bibek Raj Bhattarai, Prakriti Budhathoki, Babita Aryal, Bikash Adhikari, Ganesh Lamichhane, Darbin Kumar Poudel, Niranjan Parajuli

**Affiliations:** Central Department of Chemistry, Tribhuvan University, Kirtipur, Kathmandu, Nepal; Center for Drug Design and Molecular Simulation Division, Cancer Care Nepal and Research Center, Jorpati, Kathmandu, Nepal

**Keywords:** Natural Products, Molecular docking, Cholesterol, Enzyme inhibition

## Abstract

Hypercholesterolemia has posed a serious threat of heart diseases and stroke worldwide. Xanthine oxidase (XO), the rate-limiting enzyme in uric acid biosynthesis, is regarded as the root of reactive oxygen species (ROS) that generates atherosclerosis and cholesterol crystals. β-Hydroxy β-methylglutaryl-coenzyme A reductase (HMGR) is a rate-limiting enzyme in cholesterol biosynthesis. Although some commercially available enzyme inhibiting drugs have effectively reduced the cholesterol level, most of them have failed to meet the requirements of being apt drug candidates. Here, we have carried out an *in-silico* analysis of secondary metabolites that have already shown good inhibitory activity against XO and HMGR. Out of 118 secondary metabolites reviewed, sixteen molecules inhibiting XO and HMGR were taken based on IC_50_ values reported *in vitro* assays. Further, receptor-based virtual screening was carried out against secondary metabolites using *GOLD Protein-Ligand Docking Software*, combined with subsequent post-docking, to study the binding affinities of ligands to the enzymes*. In-Silico* ADMET analysis was carried out to study their pharmacokinetic properties, followed by toxicity prediction through ProTox-II. The molecular docking of amentoflavone **(1)** (GOLD score 70.54), and ganomycin I **(9)** (GOLD score 59.61) evinced that the drug has effectively bind at the competitive site of XO and HMGR, respectively. Besides, 6-paradol **(3)** and selgin **(4)** could be potential drug candidates to inhibit XO. Likewise, n-octadecanyl-O-α-D-glucopyranosyl(6’→1”)-O-α-D-glucopyranoside **(10)** could be potential drug candidates to maintain serum cholesterol. *In-silico* ADMET analysis showed that the sixteen metabolites were optimal within the categorical range in comparison to commercially available XO and HMGR inhibitors, respectively. Toxicity analysis through Protox-II revealed that 6-gingerol **(2)**, ganoleucoin K **(11)**, and ganoleucoin Z **(12)** are toxic for human use. This computational analysis supports earlier experimental evidence towards the inhibition of XO and HMGR by natural products. Further study is necessary to explore the clinical efficacy of these secondary molecules, which might be alternatives for the treatment of hypercholesterolemia.

**Graphical abstract:** 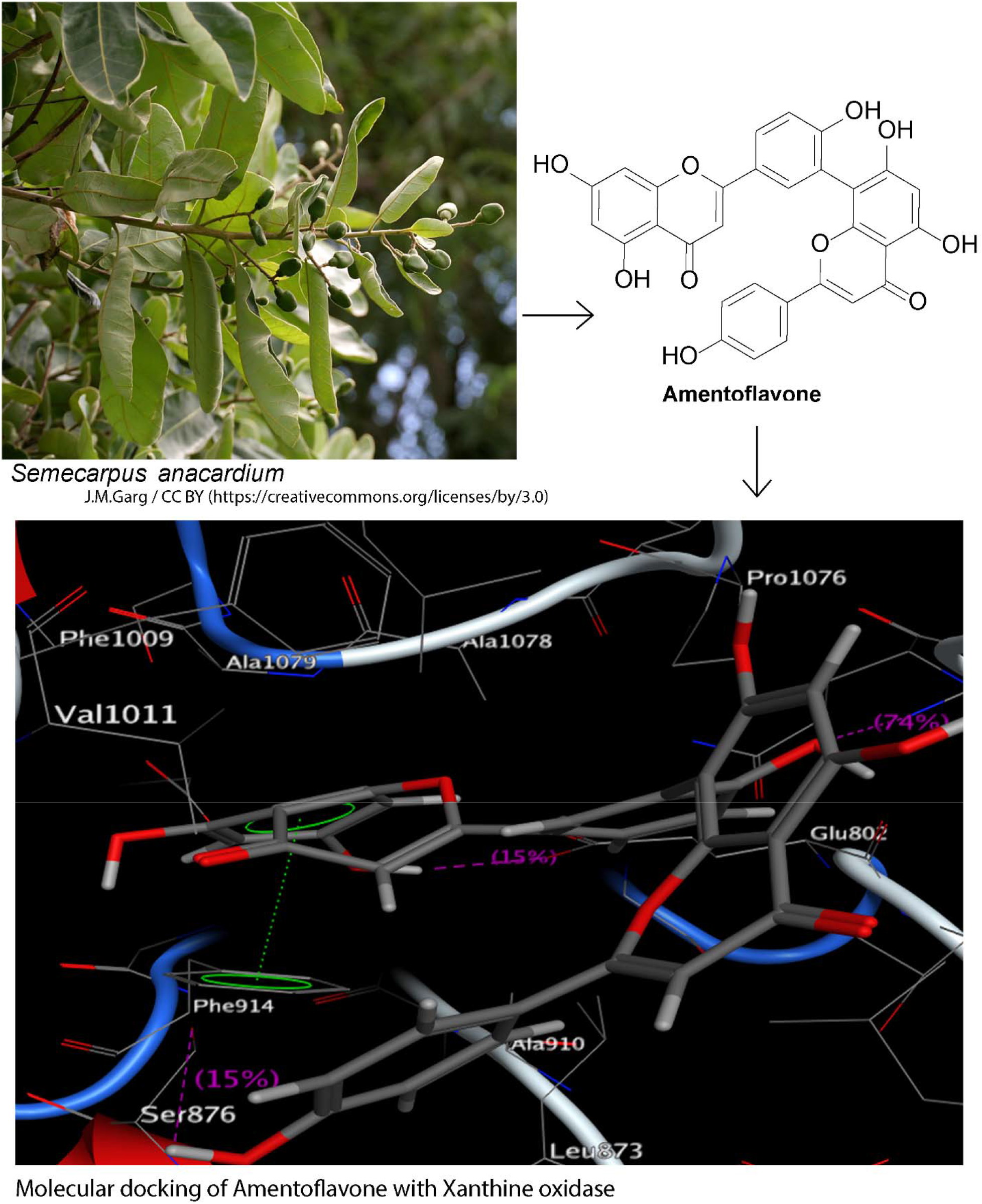

## Background

Globally, ischemic heart disease (IHD) is the foremost cause of cardiovascular diseases (CVDs) followed by stroke [1], and a high cholesterol level is accredited for one-third of IHD and increased risks of stroke [2]. Cholesterol acquired via *de novo* synthesis (600-900 mg/day) & diet (300-500 mg/day), transported via blood, and excreted through bile acid biosynthesis (500-600 mg/day) and as biliary cholesterol (600 mg/day) are the major aspects of its homeostasis in human [3], [4]. Several elements, such as age, gender, human genetics, dietary habits, physical activity, and metabolic disorder, have affected cholesterol levels [5]. Cholesterol is an indispensable structural component of the cell membrane and serves as the substrate for biosynthesis of vitamin D, bile acids, and steroid hormones [6]. However, its accumulation in the body is associated with atherosclerosis, hypertension, and ultimately to CVDs, which are found to been accountable for the increase in mortality and morbidity rates globally [1], [7]. Whence, from a therapeutic point of view, the regulations of total serum cholesterol and triglycerides have gained much heed against hyperlipidemia [8].

Statins, competitive inhibitors of HMGR, are the widely recognized medication to lower cholesterol levels [9], [10]. Other available medications include ezetimibe (cholesterol absorption and Niemann-Pick C1-like protein inhibitor) and bile acid sequestrants (induce hepatic conversion of cholesterol into bile acids) [11], [12]. Lomitapide (microsomal triglyceride transfer protein inhibitor), mipomersen (apolipoprotein B 100 inhibitor), alirocumab & evolocumab (proprotein convertase subtilisin kexin type 9 (PCSK9) inhibitors), and bempedoic acid (adenosine triphosphate-citrate lyase inhibitor) are the latest approved remedy against hypercholesterolemia in last decades [13]–[15]. Moreover, the inhibitors of acyl-CoA cholesterol acyltransferase 2 (ACAT-2), diglyceride acyltransferase 2 (DGAT-2), high-density lipoproteins (HDL) modulating drugs, small interfering RNA (inclisiran), and angiopoietin-like protein 3 (ANGPTL3) are under progress for clinical trials in human. They could be prospects in lowering cholesterol levels [10], [16].

XO (290 kDa) is involved in uric acid biosynthesis and regarded as the root of ROS (H_2_O_2_ and O_2_^−^) in vascular tissue and, hence engenders atherosclerosis [17]–[19]. It is the rate-limiting enzyme involved in the catabolism of purine nucleotides, the step that oxidizes xanthine to uric acid [20], [21]. IHD correlates with upraised levels of uric acid, and XO inhibitors such as allopurinol and febuxostat have palliated the risk of IHD by minimizing the effect of ROS and enhancing endothelial function and ATP synthesis in ischaemic tissue [22], [23]. Elevated cholesterol level increases the activity of XO that causes oxidative stress in tissue and decreases the activity of nitric oxide synthase (NOS) that surges CVDs risks [24], [25]. This oxidative stress converts low-density lipoprotein (LDL) to oxidized LDL that is absorbed by macrophages in the intima of the vascular wall that eventually forms cholesterol crystals and deteriorates endothelial function [26], [27]. In cholesterol biosynthesis, the conversion of acetyl CoA to HMG-CoA is catalyzed by HMGR (200 kDa) found in the endoplasmic reticulum (ER) [28], [29]. It is the rate-limiting enzyme involved in the synthesis of mevalonate, while the post-squalene portions are regulated by cytochrome P450 51 [3].

The mechanisms, pharmacokinetics, interactions, and side effects of the drugs, as mentioned above, are well explained by FeinGOLD [30]. On account of side effects, it is challenging to explore new drugs of high medicinal importance. In this study, we have focused on potential HMGR and XO inhibitors, based on natural products, which are considered as the wellspring of biologically and pharmacologically active sources of secondary metabolites [31], [32]. We have performed the virtual screening of some secondary metabolites showing good inhibitory activity with the aid of artificial intelligence (AI) tools. *In-silico* ADMET analysis, which would lower the use of animal testing following ethical guidelines in the pharmacological experiment, were studied using the pKCSM web application. Furthermore, toxicity analysis through ProTox-II and molecular docking using *GOLD Protein-Ligand Docking Software* combined with subsequent post-docking were carried out to uncover further evidence on the inhibition mechanism. We believe that our findings would be beneficial in drug development programs concerning hypolipidemic agents.

## Methods

### Preparation of Protein

The crystal structure of HMGR (PDB ID:1HWK, 2.22 Å) and XO (PDB ID:1N5X, 2.80 Å) were retrieved from Protein Data Bank (PDB) [33]. The dimeric crystal structure of HMGR, complexed with atorvastatin, was used for docking studies where two neighboring monomers were relevant for making interactions with statins [34] [35]. Similarly, the crystal structure of XO, complexed with febuxostat, was retrieved to understand the protein-ligand docking algorithm and to predict the position of metabolites in the binding cavity of XO. The structure of XO was a homodimer (chain A and B), where only chain A was used for the docking studies. And, other chains and water molecules were removed using the MOE protein preparation wizard [36].

### Preparation of Ligand

Sixteen plant and fungus-based secondary metabolites (**Table 1 and 2)** with potential pharmaceutical and medicinal benefits were chosen for the ligand-protein docking study. The docking study was performed against commercial drugs such as atorvastatin, simvastatin, lovastatin, and pravastatin for HMGR, prescribed to patients with high cholesterol to reduce low-density lipoprotein or bad cholesterol. On the other hand, commercial drugs such as allopurinol, febuxostat, topiroxostat, and probenecid were used for molecular docking studies with XO. The structures of the ligand molecules and the control drugs of both enzymes were retrieved from the PubChem database [37] and verified from SciFinder. The structures were retrieved in SDF format and were changed to PDB and MOL2 format using Discovery Studio Visualizer 4.0 software. The structure and complete chemical properties, torsional energy, van der Waals potential energy, electrostatic energy, weight, log *P*, total polar surface area (TPSA), donor atoms and acceptor atoms of the ligands were listed (Supplementary **Table 4S**) by the help of MOE Module [38].

**Table 1.** GOLD Fitness score and Protein-Ligand Interactions of Protein ID: 1N5X with XO Inhibitors

| Compounds | Reported IC<sub>50</sub><br>value (μM) | GOLD<br>Score | H-Bond<br>Interaction<br>Residues | Bond<br>Length | Other Interacting Residues |
| --- | --- | --- | --- | --- | --- |
| Amentoflavone (1) | 0.09 | 70.54 | Glu 802<br>Asn 768<br>Phe 914<br>Ser 876 | 2.1<br>2.7<br>-<br>2.8 | Pi-Pi interaction with Phe 914 |
| 6- Gingerol (2) | 10.50 | 68.75 | Thr 1010<br>Ala 1079<br>Phe 798<br>Phe 914 | 1.9<br>2.4<br>2.5<br>- | Pi-Pi interaction with Phe 914 |
| 6- Paradol (3) | 12.40 | 67.34 | Arg 880<br>Phe 914 | 3.2<br>- | Pi-Pi interaction with Phe 914 |
| Selgin (4) | 0.22 | 61.41 | Phe 914<br>Glu 1261<br>Thr 1010<br>Arg 880 | -<br>2.0<br>3.0<br>2.4/3.2 | Pi-Pi interaction with Phe 914 |
| Isoquercitrin (5) | 1.60 | 55.4 | Thr1010<br>Ala 1079<br>Phe 914<br>Ser 876<br>Asp 872<br>Asn 768<br>Lys 771 | 2.90/3.00<br>3.00<br>2.8<br>-<br>2.9<br>3<br>2.5 | Pi-Pi interaction with Phe 914 |
| Neotaiwanensol B (6) | 0.28 | 52.77 | Ser 876<br>Thr 1010<br>Arg 880<br>Phe 914 | 3<br>1.6<br>2.90/2.90<br>2.9 | Pi-Pi interaction with Phe 914 |
| Hydroxychavicol (7) | 0.38 | 49.02 | Arg 880<br>Phe 914 | 2.9/2.7<br>- | Pi-Pi interaction with Phe 914 |
| Riparsaponin (8) | 0.01 | 27.92 | Leu 648 | 3.2 | No any extra interaction |

**Table 2.** GOLD Fitness score and Protein-Ligand Interactions of Protein ID: 1HWK with HMGR Inhibitors.

| Compounds | Reported IC<sub>50</sub> value (μM) | GOLD Score | H-Bond Interaction Residues | Bond Length (Å) | Other Interacting Residues |
| --- | --- | --- | --- | --- | --- |
| Ganomycin I (**9**) | 12.3 ± 1.7 | 59.61 | Arg A590 | - | Pi-cation interaction with Arg A590 |
| n-octadecanyl-O-α-D-glucopyranosyl(6'→1'')-O-α-D-glucopyranoside (**10**) | 0.164 | 52.69 | Lys A691<br>Gly B560<br>Lys B735<br>Arg A590<br>Asp A690 | 3.1<br>2.8<br>3.2<br>2.9<br>1.7/3.3 | No any extra interaction |
| Ganoleucoin K (**11**) | 10.7 ± 2.9 | 43.91 | Asn A658<br>Val A805 | 2.5<br>2.4 | No any extra interaction |
| Ganoleucoin Z (**12**) | 8.68 ± 0.96 | 43.41 | Lys A691<br>Asp A690<br>Arg A590<br>Lys B735<br>Leu B857 | 2.8<br>2.10/ 3.30<br>2.7<br>3.1<br>2.2 | No any extra interaction |
| Ganoleucoin Y (**13**) | 9.72 ± 0.91 | 30.85 | Asn A658<br>Ser A661 | 2.6<br>2.3 | No any extra interaction |
| Ganoleucoin T (**14**) | 10.3 ± 1.78 | 29.15 | Asp A690<br>Gly B560<br>Thr B558 | 1.3<br>3.2<br>2.2 | No any extra interaction |
| Ganoderic acid DM (**15**) | 9.5 ± 1.5 | 18.89 | Arg A590<br>Asn B755 | 2.7<br>3.0 | No any extra interaction |
| Ganoderic acid $\eta$ (**16**) | 29.8 ± 1.5 | 12.13 | Arg A590<br>Lys A692<br>Lys B735 | 2.5<br>2.9<br>2.8 | No any extra interaction |

### Prediction of active sites

Amino acids involved in active pocket formation were determined using Site-Finder (Supplementary **Table 3S**), which calculates possible active sites in a receptor from the 3D atomic coordinates based on alpha shape methodology [39]. From the active site analysis, all the amino acid residues were listed adequately and validated from published crystal structure active residues along with published research journals for complete study [40], [41].

### Computational analysis

Flexible docking simulations were performed using GOLD [42] to investigate binding modes of the molecules to predict the efficiency of secondary metabolites for the inhibition of HMGR and XO enzymes. These novel potential compounds were obtained from the extensive literature review and deposited in the inbuilt CHEM-TU Natural Metabolites Library. GOLD uses a genetic algorithm for docking and performs automated docking with fully ligand flexibility and partial flexibility in the neighborhood of the protein active site [43] to determine the appropriate binding positions, orientations, and conformations of ligands [44]. All other parameters were maintained as default. In the flexible docking process, the compound which has the highest fitness score was considered to have the highest binding affinity according to GOLD score molecular mechanics function which is the combination of various parameters expressed as,

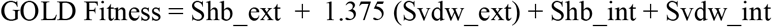

Where Shb_ext is the protein-ligand hydrogen-bond score, and Svdw_ext is the protein-ligand van der Waals score. Shb_int is the contribution to fitness due to intramolecular hydrogen bonds in the ligand, Svdw_int is the contribution due to intermolecular strain in the ligand. The details about molecular docking results are mentioned in Supplementary **Table 1S**.

### Prediction of ADMET profiles

Drug discovery programs assisted analysis of absorption, distribution, metabolism, excretion, and toxicity (ADMET) properties of secondary metabolites. The potential pharmacokinetic properties prediction was completed using the pKCSM web application [45]. *In-silico* potential toxicity of secondary metabolites was assessed by ProTox-II, which was based on toxic, lethal dose (LD_50_) value ranging from class 1 and 2 (fatal), class 3 (toxic), class 4 and 5 (harmful), while class-6 (non-toxic) [46]. The confidence score of secondary metabolites for specific targets had been used to predict the reliability of toxicity based on the value of more than 0.7 [47].

## Results

In the beginning, a dataset was prepared based on a literature review taking IC_50_ values of *in-vitro* enzyme inhibition assays with HMGR and XO by natural products. In this article, Supplementary **Table 2S** provides details about the targets and their description. **Figure 1** provides the structure of secondary metabolites and natural sources, which were used in this study. **Tables 1 and 2** give GOLD fitness scores and hydrogen bonding interaction values between targets and secondary metabolites, interaction type, and bond length of the docking. The 2D and 3D interactions of the high GOLD scoring metabolites and commercial drugs with the target enzymes were shown in **Figure 2-3,** and Supplementary **Figure (1S-7S)**. The molecular properties of commercial drugs and selected secondary metabolites were shown in Supplementary **Table 4S**.

**Figure 1.**
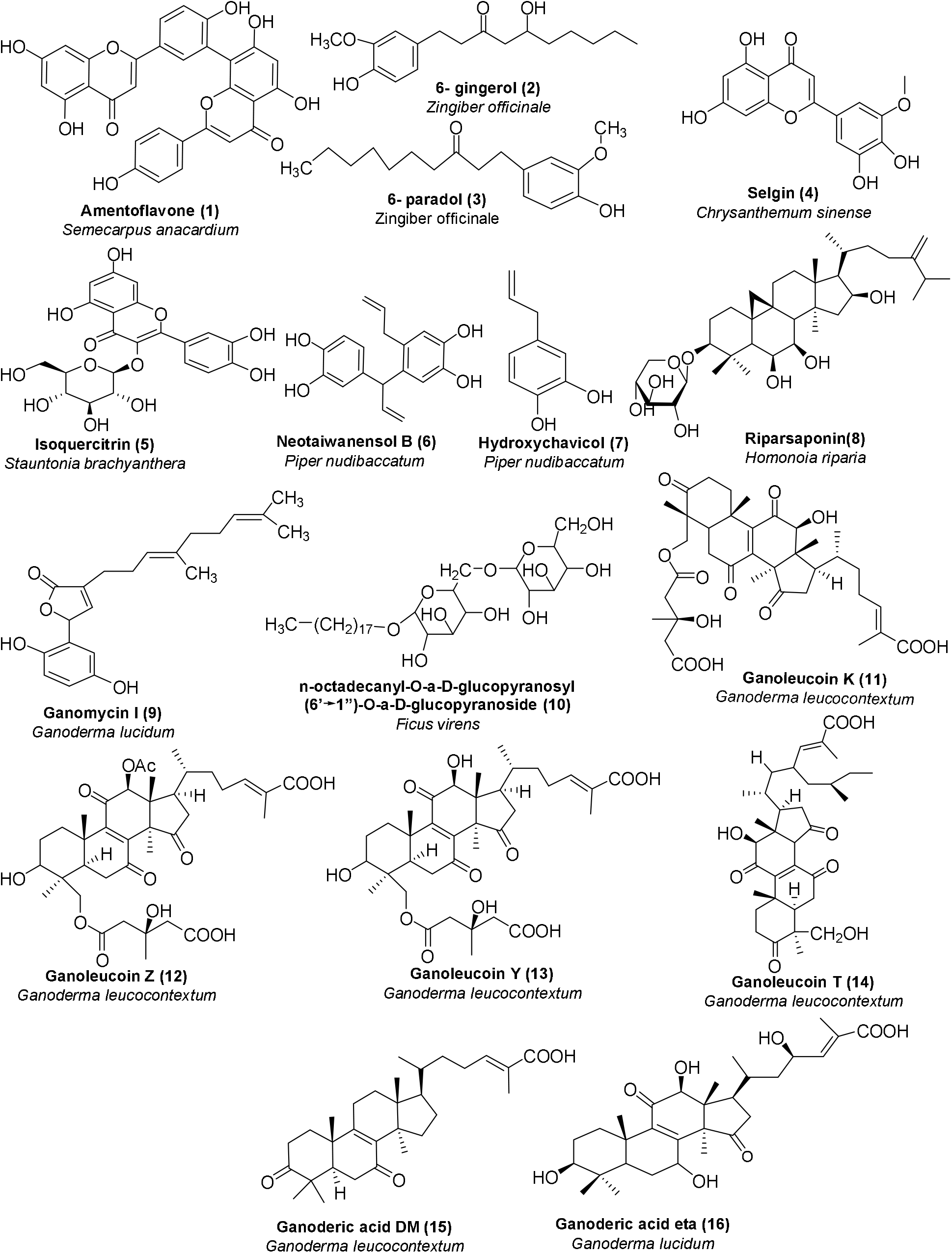
Plant and fungus-based secondary metabolites inhibiting XO (**1-8**) and HMGR (**9-16**)

**Figure 2.**
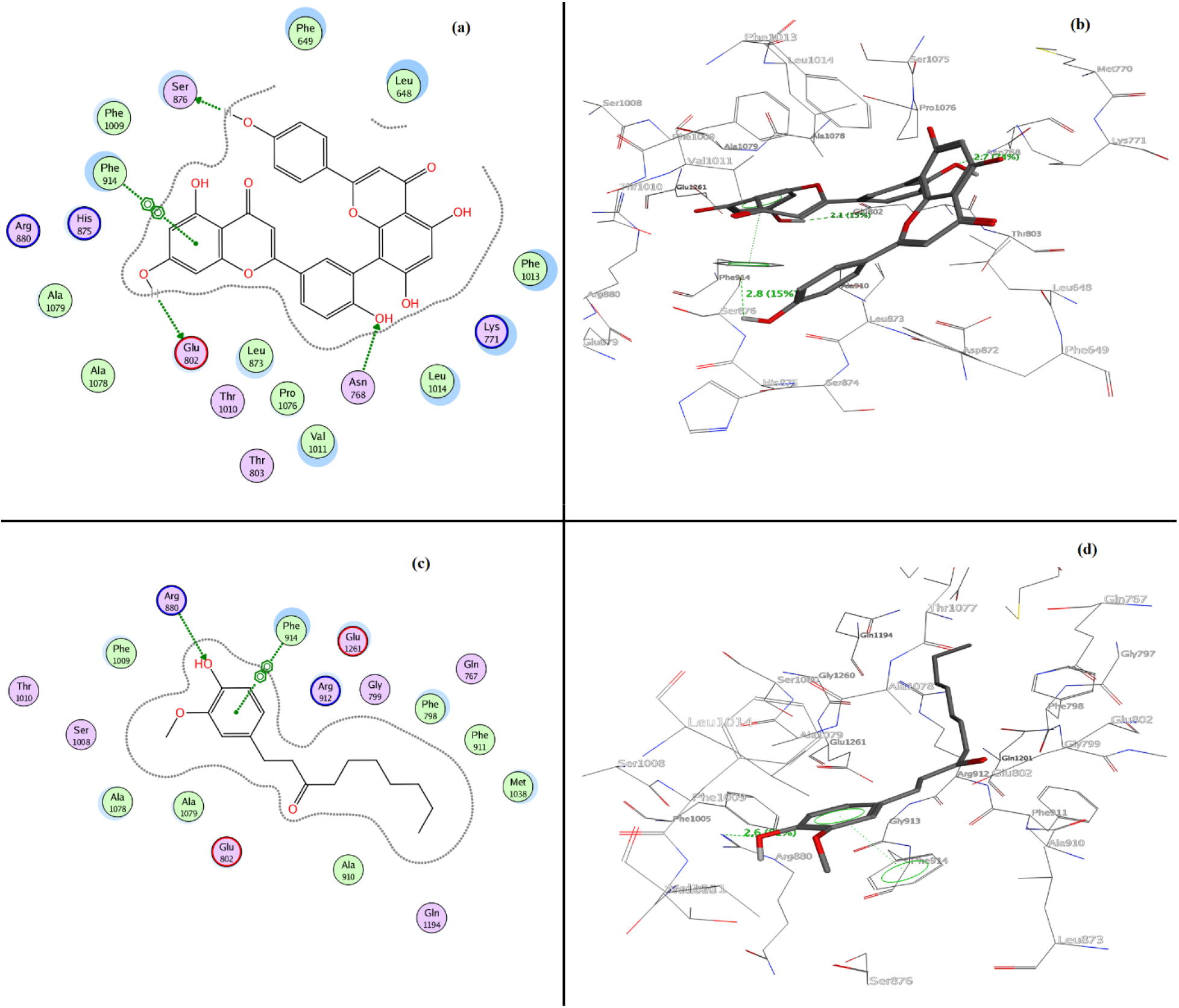
(**a**) 2D and (**b**) 3D interactions of XO with amentoflavone **(1**) (GOLD fitness score of 70.54); (**c**) 2D and (**d**) 3D (lower) interactions of XO with 6-gingerol **(2)** (GOLD fitness score of 68.75)

**Figure 3.**
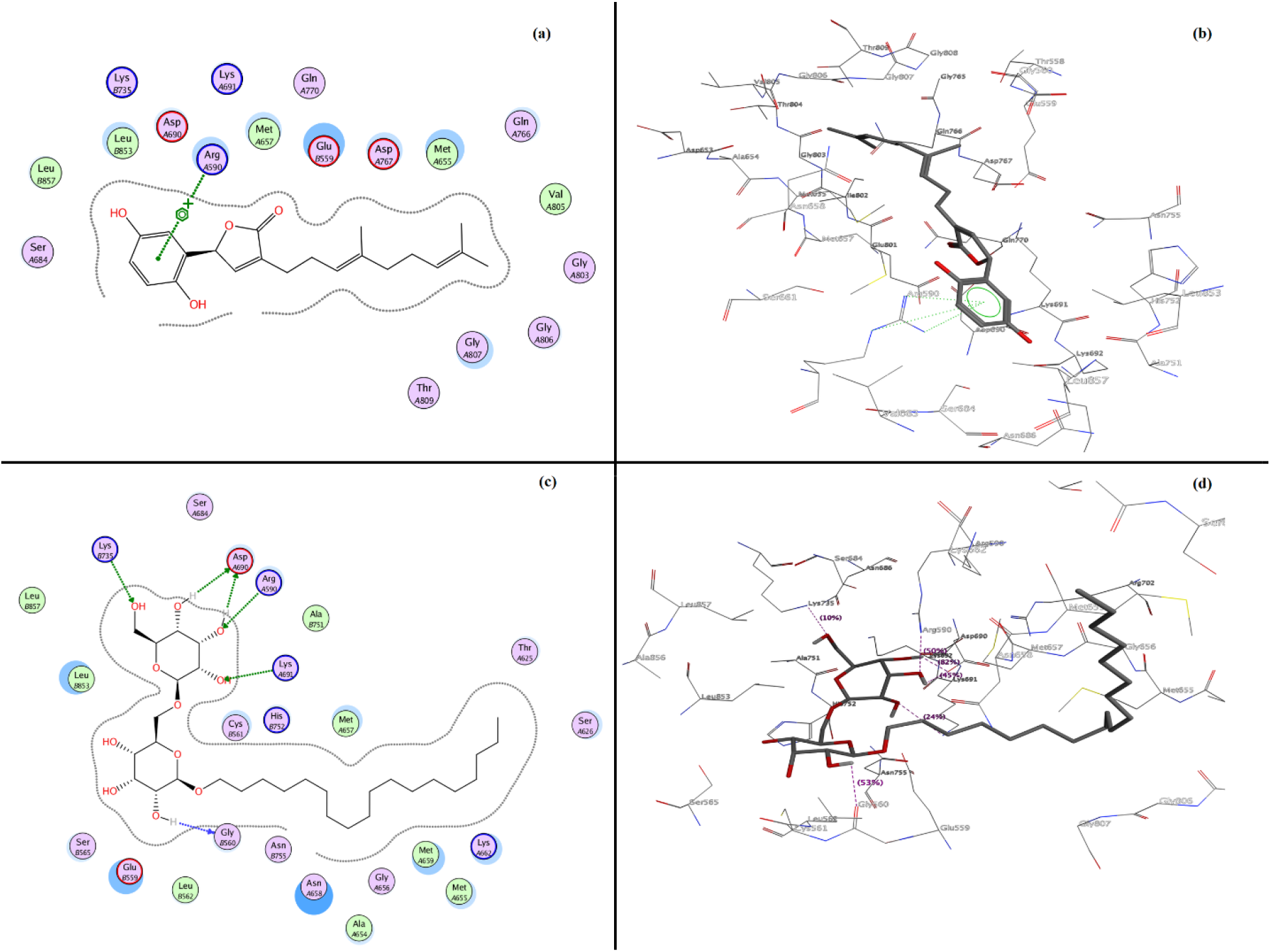
(**a**) 2D and (**b**) 3D interactions of HMGR with ganomycin-I **(9)** (GOLD fitness score of 59.61); (**c**) 2D and (**d**) 3D interactions of HMGR with n-octadecanyl-O- -D-glucopyranosyl(6’→ 1”)-O- -D-glucopyranoside **(10**) (GOLD fitness score of 52.69)

### XO Molecular Docking Analysis

From the molecular docking, we observed electrostatic 2D and 3D molecular surfaces (**Figure 2)**, the study showed that amentoflavone **(1)** and 6-paradol **(3)** were well located into the active site of XO with the GOLD fitness score of 70.54, and 67.34 **(Table 1)** respectively which is higher than commercial drugs febuxostat (GOLD score 64.53), topiroxostat (GOLD score 61.46), probenecid (GOLD score 57.75), and allopurinol (GOLD score 46.16) (Supplementary **Table 10S**). Similarly, selgin **(4)**, isoquercitrin **(5)**, neotaiwanesol B **(6)**, and hydroxychavicol **(7)** have shown satisfactory interactions with the active site residues with fitness score of 61.41, 55.4, 52.77, and 49.02 respectively.

Amino acid residues participating in forming Pi-Pi and Pi-cation interactions were also investigated. Amentoflavone **(1)** and 6-paradol **(3)** was surrounded by several amino acid residues (Glu 802, Asn 768, Phe 914, ser 876, and Arg 880), which were described as active site residues [48]. It has been reported that Arg 880 and Glu 802 residues play a key role in the hydroxylation of substrate xanthine [49]. Hydrogen bonding with Glu 802 residue for a top-scoring amentoflavone **(1)** and Arg 880 for commercial drugs (febuxostat) was observed with the bond lengths of 2.1 Å and 3.0 Å, respectively. Nevertheless, 6-paradol **(3)**, selgin **(4)**, and isoquercitrin **(5)** also showed significant interactions through H-bonding within the range of 2-3.2 Å as well as Pi-Pi interaction with Phe 914 of target XO protein.

### HMGR Molecular Docking Analysis

**Figure 3** showed the 2D and 3D molecular surface’s orientation of the top-scored secondary metabolites ganomycin I **(9)**, and n-octadecanyl-O-α-D-glucopyranosyl(6’→1”)-O-α-D-glucopyranoside **(10)**. The study showed that ganomycin I **(9)** bound competitively into the active site of HMGR with the GOLD fitness score of 59.61 **(Table 2)** higher than commercial drugs simvastatin (GOLD fitness score 56.81), lovastatin (GOLD fitness score 41.36) and pravastatin (GOLD fitness score 54.83) (Supplementary **Table 10S).** Similarly, n-octadecanyl-O-α-D-glucopyranosyl(6’→1”)-O-α-D-glucopyranoside **(10)** has shown satisfactory interactions with the active site residues with fitness scores of 52.69. Arg A590 amino acid was found to be involved in forming Pi-cation interaction with ganomycin I **(9)** having unique features of high Vander Waals energy 32.11 (Supplementary **Table 4S**). In commercial drugs, other active residues (Arg A590, Ser A684, Gly A692, Lys A691, Asp A690, Glu B559, Lys B735) were involved in forming hydrogen bonds as wells as in Pi-Pi interactions. Furthermore, ganoleucoin T **(14)**, ganoderic acid DM **(15)**, and ganoderic acid η **(16)** were found interacting with target protein amino acid residues via H-bonds ranging 1.3-3.3 Å (**Figure 3**).

### Analysis of ADMET profiles

Supplementary **Table 8S (a)** and **8S (b)** showed the detail of the ADMET analysis of sixteen metabolites. Compounds (**1) – (9)** and (**14) – (16)** were significantly absorbed, while (**10)** – (**13)** were found to be poorly absorbed in the human small intestine. However, these metabolites were unable to cross blood-brain barriers (BBB) readily, and none of this inhibited CYP3A4 and CYP2D6. Utterly, all the metabolites showed low hepatic and renal clearance. According to the *in-silico* toxicity prediction through Protox-II, compounds (**14**) and (**16**) were non-toxic while compounds (**3**), (**4**), (**5**), (**8**), (**1**), (**6**), (**7**), (**9**), (**10**), and (**15**) were classified under the harmful category. Compounds (**2**), (**11**), (**12**), and (**13**) were found to be toxic for human use. Moreover, Supplementary **Table 7S** showed the detail of predicted LD_50_ values and confidence scores of specific active targets of each metabolite. This analysis showed that all sixteen metabolites except (**2**), (**11**), (**12**), and (**13**) were optimal within the categorical range compared to commercially available XO and HMGR inhibitors, respectively.

## Discussion

Active plant-derived drugs can create a new era as therapeutic agents. Although this article is based on molecular docking of natural products already characterized, **Table 9S** shows crude extracts of natural sources that show significant inhibitory activity against XO and HMGR for broader coverage in the field. Molecular docking continues to hold great promise in the field of computer-based drug design. Higher the GOLD fitness score of ligands, the higher would be the binding capacity of it to protein residues [44]. Hydrogen bonding and hydrophobic interactions play an important role in determining the binding affinity and stability of protein-ligand complexes [50], [51].

The GOLD fitness scoring of **(1)** and **(3)** was found to be higher than commercially available XO inhibitors. Moreover, the GOLD fitness score of (**4**), **(5)**, **(6)**, and **(7)** are harmonious with that of commercial XO inhibitors. For instance, the inhibitory effect of the XO is probably due to lodging of **(1)** in its active site via H-bonding with neighboring Glu 802, Asn 768, Ser 876, and Phe 914 amino acid residues and Pi-Pi interaction with Phe 914. From this interaction, the catalytic center of XO would undergo a conformational change and suppress the activity of the enzyme [52]. XO is a complex molybdoflavoprotein that produces ROS via the reduction of oxygen at the flavin center [53]. With the inhibition of XO, increased levels of HDL have been reported in animal models [54]. Moreover, HDL is associated with reverse cholesterol transport and cholesterol efflux that prevents atherosclerosis and, ultimately, CVDs [55]. Moreover, the studies showed the superiority of non-purine analog of XO inhibitors over purine analogs in terms of side effects and toxicity [56]. On account of this, the compounds **1-8** are non-purine analogs. Previously, **(1)** and **(3)** were shown with anticancer, anti-inflammatory, and antioxidant activities [57]–[60]. Thus, this is evident that XO inhibitors, used widely as the remedy of hyperuricemia, could be potential candidates for regulating body cholesterol levels.

Similarly, the binding affinity of **(9)** to enzyme active sites, via H-bonding and Pi-Pi interaction with Arg A590 amino acid residue, is found to be higher than that of commercial inhibitors of HMGR. The probable mechanism is competitive inhibition via the binding of **(9)** and **(10)** at the active site of HMGR inhibiting natural substrate (HMG-CoA). This inhibition lowers the cholesterol levels in the ER and induces the transport of sterol regulatory element-binding proteins (SREBPs) to the Golgi body. These SREBPs express the genes of LDL receptors and speed up the removal of LDL and VLDL from plasma [30], [61]. Previously, **(9)** had been shown as HMGR as well as an α-glucosidase inhibitor [62]. This showed that these secondary metabolites also bind effectively to enzymes like commercial drugs and could be potential inhibitors of XO and HMGR, respectively. Though the compounds **(2)** and **(11)-(13)** possess appropriate GOLD scores, they are not regarded as potential drugs owing to their toxicity on virtual screening.

In ADMET profiles, intestinal absorption value above 30% signifies good absorption in the human intestine. The volume of distribution (VDss) is taken into consideration if logVDss value greater than 0.45. The compounds with logBB < −1 are said to be poorly distributed to brain, while those having logBB >0.3 are potential to cross BBB [45], [63], [64]. The cytochrome P450 (CYP) plays a major role in drug metabolism with CYP (1A2, 2C9, 2C19, 2D6, and 3A4), mainly responsible for the biotransformation of greater than 90% of drugs in phase-1 metabolism [65], [66]. However, among these P450 families, CYP3A4 is the focus part of this study [67]. The relationship between the rate of elimination of the drug and concentration of the drug in the body is best described by total clearance [68]. Moreover, it is mandatory to examine the toxicity value based on parameters like ames toxicity, hepatotoxicity, and oral toxicity range because these play a critical role in the selection of drugs.

As discussed earlier, the studies suggest that XO inhibitors lessen the threat of major adverse cardiovascular events (MACE) [69]. With the availability of a handful of XO inhibitors in clinical use for treating hyperuricemia, the novel drugs addressing cardiac complications are demanding [70]. Recently, non-purine-like XO inhibitors have drawn significant attention, so they do not interfere with other facets of purine metabolism [48]. XO inhibitors could be equally useful for diabetic patients [71]. Although statins have been used widely for many decades as HMGR inhibitors, they are marked with muscular disorders, diabetes, liver diseases, and so on [30], [72]. Novel HMGR inhibitors are called for minimizing these side effects. However, the problems associated with natural secondary metabolites such as their stability, solubility, and bioavailability need to be taken into account to use them as therapeutic agents [73]. Furthermore, clinical trials, structural modifications, and biomimetic synthesis may lead to the discovery of promising inhibitors of HMGR and XO, which could contribute to the treatment of hyperuricemia and lowering cholesterol.

## Conclusion

The most potent natural inhibitors of the XO and HMGR are selected based on reported IC_50_ values and show that these metabolites could be potent in lowering cholesterol levels. The molecular docking analysis shows that amentoflavone **(1)**, 6-paradol **(3)**, and selgin **(4)** fit well in the binding site of XO and lower the catalytic activity of the enzyme by changing its conformation. The studies and evidence suggest that XO inhibitors could be potential in regulating cholesterol levels. Similarly, the binding affinity of ganomycin-I **(9)** and n-octadecanyl-O-α-D-glucopyranosyl(6’→1”)-O-α-D-glucopyranoside **(10)** are involved in arresting the cholesterol biosynthetic pathway. Thus, the shreds of evidence like experimental IC_50_ value, computational docking, *in-silico* pharmacokinetics, and toxicity analysis proclaim that natural products could be used to develop potential future drug candidates.

## List of abbreviations

ACAT-2: Acyl-CoA cholesterol acyltransferase 2
AI: Artificial intelligence
ANGPTL3: Angiopoietin-like protein 3
ATP: Adenosine triphosphate
BBB: Blood-brain barrier
CVDs: Cardiovascular diseases
CYP: Cytochrome P450
DGAT-2: Diglyceride acyltransferase 2
ER: Endoplasmic reticulum
GOLD: Genetic optimization for ligand docking
HDL: High-density lipoproteins
HMG-CoA: β-Hydroxy β-methylglutaryl-coenzyme A
HMGR: HMG-CoA reductase
IHD: Ischemic heart disease
kDa: Kilodalton
LDL: Low-density lipoprotein
MACE: Major adverse cardiovascular events
MOE: Molecular Operating Environment
NOS: Nitric oxide synthase
PCSK9: Proprotein convertase subtilisin kexin type 9
ROS: Reactive oxygen species
SREBPs: Sterol regulatory element-binding proteins
TC: Total cholesterol
TG: Triglyceride
TPSA: Total polar surface area
VLDL: Very low-density lipoprotein
WHO: World Health Organization
XO: Xanthine oxidase

## Declarations

### Ethics approval and consent to participate

Not applicable

### Consent for publication

The authors declare that there is no conflict of interest regarding the publication of this paper.

### Availability of data and materials

The datasets used and/or analyzed during the current study are available from the corresponding author on reasonable request.

### Competing interests

The authors declare that they have no competing interests

### Funding

No

### Authors’ contributions

RM, BRB, and PB analyzed the literature on XO and HMGR and interpreted data. SB performed the molecular docking experiments and was a major contributor in writing the manuscript. BA, GL, BRB, DKP, BA, and NP drafted the manuscript. NP supervised the project and developed the concept. All authors have read and approved the final manuscript.

## Acknowledgements

We thank Mr. Sita Ram Phuyal, Mr. Purushottam Niraula, Mr. Karan Khadayat, and Mrs. Sonika Dawadi for constructive suggestions.

## Notes

### Competing Interest Statement

The authors have declared no competing interest.

## References

[1] P. Joseph et al., “Reducing the Global Burden of Cardiovascular Disease, Part 1: The Epidemiology and Risk Factors,” Circ. Res., vol. 121, no. 6, pp. 677–694, Sep. 2017, doi: 10.1161/CIRCRESAHA.117.308903.

[2] “WHO,” WHO, Raised Cholesterol. https://www.who.int/gho/ncd/risk_factors/cholesterol_text/en/ (accessed Jul. 05, 2020).

[3] I. A. Pikuleva, “Cholesterol-metabolizing cytochromes P450: implications for cholesterol lowering,” Expert Opin. Drug Metab. Toxicol., vol. 4, no. 11, pp. 1403–1414, Nov. 2008, doi: 10.1517/17425255.4.11.1403.

[4] J. J. Repa and D. J. Mangelsdorf, “The Role of Orphan Nuclear Receptors in the Regulation of Cholesterol Homeostasis,” Annu. Rev. Cell Dev. Biol., vol. 16, no. 1, pp. 459–481, Nov. 2000, doi: 10.1146/annurev.cellbio.16.1.459.

[5] D. Weissglas-Volkov and P. Pajukanta, “Genetic causes of high and low serum HDL-cholesterol,” J. Lipid Res., vol. 51, no. 8, pp. 2032–2057, Aug. 2010, doi: 10.1194/jlr.R004739.

[6] J.-P. Liu, Y. Tang, S. Zhou, B. H. Toh, C. McLean, and H. Li, “Cholesterol involvement in the pathogenesis of neurodegenerative diseases,” Mol. Cell. Neurosci., vol. 43, no. 1, pp. 33–42, Jan. 2010, doi: 10.1016/j.mcn.2009.07.013.

[7] X.-H. Yu, D.-W. Zhang, X.-L. Zheng, and C.-K. Tang, “Cholesterol transport system: An integrated cholesterol transport model involved in atherosclerosis,” Prog. Lipid Res., vol. 73, pp. 65–91, Jan. 2019, doi: 10.1016/j.plipres.2018.12.002.

[8] S. S. Alvi, D. Iqbal, S. Ahmad, and M. S. Khan, “Molecular rationale delineating the role of lycopene as a potent HMG-CoA reductase inhibitor: in vitro and in silico study,” Nat. Prod. Res., vol. 30, no. 18, pp. 2111–2114, Sep. 2016, doi: 10.1080/14786419.2015.1108977.

[9] R. S. Blumenthal, “Statins: Effective antiatherosclerotic therapy,” Am. Heart J., vol. 139, no. 4, pp. 577–583, Apr. 2000, doi: 10.1016/S0002-8703(00)90033-4.

[10] A. Pirillo, G. D. Norata, and A. L. Catapano, “LDL-Cholesterol-Lowering Therapy,” Berlin, Heidelberg: Springer Berlin Heidelberg, 2020.

[11] H. E. Bays, D. Neff, J. E. Tomassini, and A. M. Tershakovec, “Ezetimibe: cholesterol lowering and beyond,” Expert Rev. Cardiovasc. Ther., vol. 6, no. 4, pp. 447–470, Apr. 2008, doi: 10.1586/14779072.6.4.447.

[12] M. Mazidi, P. Rezaie, E. Karimi, and A. P. Kengne, “The effects of bile acid sequestrants on lipid profile and blood glucose concentrations: A systematic review and meta-analysis of randomized controlled trials,” Int. J. Cardiol., vol. 227, pp. 850–857, Jan. 2017, doi: 10.1016/j.ijcard.2016.10.011.

[13] R. P. Giugliano and M. S. Sabatine, “Are PCSK9 Inhibitors the Next Breakthrough in the Cardiovascular Field?,” J. Am. Coll. Cardiol., vol. 65, no. 24, pp. 2638–2651, Jun. 2015, doi: 10.1016/j.jacc.2015.05.001.

[14] U. Laufs et al., “Efficacy and Safety of Bempedoic Acid in Patients With Hypercholesterolemia and Statin Intolerance,” J. Am. Heart Assoc., vol. 8, no. 7, Apr. 2019, doi: 10.1161/JAHA.118.011662.

[15] D. J. Rader and J. J. P. Kastelein, “Lomitapide and Mipomersen: Two First-in-Class Drugs for Reducing Low-Density Lipoprotein Cholesterol in Patients With Homozygous Familial Hypercholesterolemia,” Circulation, vol. 129, no. 9, pp. 1022–1032, Mar. 2014, doi: 10.1161/CIRCULATIONAHA.113.001292.

[16] M. W. Huff, A. Daugherty, and H. Lu, “Atherosclerosis,” in Biochemistry of Lipids, Lipoproteins and Membranes, Elsevier, 2016, pp. 519–548.

[17] N. Aziz and R. T. Jamil, “Biochemistry, Xanthine Oxidase,” in StatPearls, Treasure Island (FL): StatPearls Publishing, 2020.

[18] J. R. Murrow, “Statins, Diabetic Oxidative Stress and Vascular Tissue,” in Diabetes: Oxidative Stress and Dietary Antioxidants, Elsevier, 2014, pp. 183–190.

[19] P. Pacher, S. Bátkai, and G. Kunos, “The Endocannabinoid System as an Emerging Target of Pharmacotherapy,” Pharmacol. Rev., vol. 58, no. 3, pp. 389–462, Sep. 2006, doi: 10.1124/pr.58.3.2.

[20] M. G. Battelli, A. Bolognesi, and L. Polito, “Pathophysiology of circulating xanthine oxidoreductase: New emerging roles for a multi-tasking enzyme,” Biochim. Biophys. Acta BBA – Mol. Basis Dis., vol. 1842, no. 9, pp. 1502–1517, Sep. 2014, doi: 10.1016/j.bbadis.2014.05.022.

[21] J. Maiuolo, F. Oppedisano, S. Gratteri, C. Muscoli, and V. Mollace, “Regulation of uric acid metabolism and excretion,” Int. J. Cardiol., vol. 213, pp. 8–14, Jun. 2016, doi: 10.1016/j.ijcard.2015.08.109.

[22] A. Struthers and F. Shearer, “Allopurinol: novel indications in cardiovascular disease,” Heart, vol. 98, no. 21, pp. 1543–1545, Nov. 2012, doi: 10.1136/heartjnl-2012-302249.

[23] M. Zdrenghea, A. Sitar-Tǎut, G. Cismaru, D. Zdrenghea, and D. Pop, “Xanthine oxidase inhibitors in ischaemic heart disease,” Cardiovasc. J. Afr., vol. 28, no. 3, pp. 201–204, Jun. 2017, doi: 10.5830/CVJA-2016-068.

[24] E. Devrim, İ. B. Ergüder, H. Özbek, and İ. Durak, “High-cholesterol diet increases xanthine oxidase and decreases nitric oxide synthase activities in erythrocytes from rats,” Nutr. Res., vol. 28, no. 3, pp. 212–215, Mar. 2008, doi: 10.1016/j.nutres.2008.01.006.

[25] J. Saban-Ruiz, A. Alonso-Pacho, M. Fabregate-Fuente, and C. de la Puerta Gonzalez-Quevedo, “Xanthine Oxidase Inhibitor Febuxostat as a Novel Agent Postulated to Act Against Vascular Inflammation,” Anti-Inflamm. Anti-Allergy Agents Med. Chem., vol. 12, no. 1, pp. 94–99, Jan. 2013, doi: 10.2174/1871523011312010011.

[26] G. K. Hansson and A. Hermansson, “The immune system in atherosclerosis,” Nat. Immunol., vol. 12, no. 3, pp. 204–212, Mar. 2011, doi: 10.1038/ni.2001.

[27] M. Mudau, A. Genis, A. Lochner, and H. Strijdom, “Endothelial dysfunctionL: the early predictor of atherosclerosis,” Cardiovasc. J. Afr., vol. 23, no. 4, pp. 222–231, May 2012, doi: 10.5830/CVJA-2011-068.

[28] E. S. Istvan, “Structural Mechanism for Statin Inhibition of HMG-CoA Reductase,” Science, vol. 292, no. 5519, pp. 1160–1164, May 2001, doi: 10.1126/science.1059344.

[29] H. M. Miziorko, “Enzymes of the mevalonate pathway of isoprenoid biosynthesis,” Arch. Biochem. Biophys., vol. 505, no. 2, pp. 131–143, Jan. 2011, doi: 10.1016/j.abb.2010.09.028.

[30] K. R. Feingold, “Cholesterol Lowering Drugs,” in Endotext, South Dartmouth (MA): MDText.com, Inc., 2020.

[31] A. G. Atanasov et al., “Discovery and resupply of pharmacologically active plant-derived natural products: A review,” Biotechnol. Adv., vol. 33, no. 8, pp. 1582–1614, Dec. 2015, doi: 10.1016/j.biotechadv.2015.08.001.

[32] T. Linder et al., “Design and Synthesis of a Compound Library Exploiting 5-Methoxyleoligin as Potential Cholesterol Efflux Promoter,” Molecules, vol. 25, no. 3, p. 662, Feb. 2020, doi: 10.3390/molecules25030662.

[33] S. K. Burley et al., “RCSB Protein Data Bank: Sustaining a living digital data resource that enables breakthroughs in scientific research and biomedical education: RCSB Protein Data Bank,” Protein Sci., vol. 27, no. 1, pp. 316–330, Jan. 2018, doi: 10.1002/pro.3331.

[34] K. Okamoto, B. T. Eger, T. Nishino, S. Kondo, E. F. Pai, and T. Nishino, “An Extremely Potent Inhibitor of Xanthine Oxidoreductase: CRYSTAL STRUCTURE OF THE ENZYME-INHIBITOR COMPLEX AND MECHANISM OF INHIBITION,” J. Biol. Chem., vol. 278, no. 3, pp. 1848–1855, Jan. 2003, doi: 10.1074/jbc.M208307200.

[35] S.-Y. Jiang et al., “Discovery of a potent HMG-CoA reductase degrader that eliminates statin-induced reductase accumulation and lowers cholesterol,” Nat. Commun., vol. 9, no. 1, p. 5138, Dec. 2018, doi: 10.1038/s41467-018-07590-3.

[36] M. D. Santi et al., “Xanthine oxidase inhibitory activity of natural and hemisynthetic flavonoids from Gardenia oudiepe (Rubiaceae) in vitro and molecular docking studies,” Eur. J. Med. Chem., vol. 143, pp. 577–582, Jan. 2018, doi: 10.1016/j.ejmech.2017.11.071.

[37] E.-K. Kwon et al., “Flavonoids from the Buds of Rosa damascena Inhibit the Activity of 3-Hydroxy-3-methylglutaryl-coenzyme A Reductase and Angiotensin I-Converting Enzyme,” J. Agric. Food Chem., vol. 58, no. 2, pp. 882–886, Jan. 2010, doi: 10.1021/jf903515f.

[38] R. Guha and E. Willighagen, “A Survey of Quantitative Descriptions of Molecular Structure,” Curr. Top. Med. Chem., vol. 12, no. 18, pp. 1946–1956, Sep. 2012, doi: 10.2174/156802612804910278.

[39] C. A. Del Carpio, Y. Takahashi, and S. Sasaki, “A new approach to the automatic identification of candidates for ligand receptor sites in proteins: (I) Search for pocket regions,” J. Mol. Graph., vol. 11, no. 1, pp. 23–29, Mar. 1993, doi: 10.1016/0263-7855(93)85003-9.

[40] J. Zhang et al., “Eight new triterpenoids with inhibitory activity against HMG-CoA reductase from the medical mushroom Ganoderma leucocontextum collected in Tibetan plateau,” Fitoterapia, vol. 130, pp. 79–88, Oct. 2018, doi: 10.1016/j.fitote.2018.08.009.

[41] V. B. da Silva, C. A. Taft, and C. H. T. P. Silva, “Use of Virtual Screening, Flexible Docking, and Molecular Interaction Fields To Design Novel HMG-CoA Reductase Inhibitors for the Treatment of Hypercholesterolemia ^†^,” J. Phys. Chem. A, vol. 112, no. 10, pp. 2007–2011, Mar. 2008, doi: 10.1021/jp075502e.

[42] G. Jones, P. Willett, R. C. Glen, A. R. Leach, and R. Taylor, “Development and validation of a genetic algorithm for flexible docking,” J. Mol. Biol., vol. 267, no. 3, pp. 727–748, Apr. 1997, doi: 10.1006/jmbi.1996.0897.

[43] V. Z. Spassov and L. Yan, “A fast and accurate computational approach to protein ionization,” Protein Sci., vol. 17, no. 11, pp. 1955–1970, Nov. 2008, doi: 10.1110/ps.036335.108.

[44] N. S. Pagadala, K. Syed, and J. Tuszynski, “Software for molecular docking: a review,” Biophys. Rev., vol. 9, no. 2, pp. 91–102, Apr. 2017, doi: 10.1007/s12551-016-0247-1.

[45] D. E. V. Pires, T. L. Blundell, and D. B. Ascher, “pkCSM: Predicting Small-Molecule Pharmacokinetic and Toxicity Properties Using Graph-Based Signatures,” J. Med. Chem., vol. 58, no. 9, pp. 4066–4072, May 2015, doi: 10.1021/acs.jmedchem.5b00104.

[46] P. Banerjee, A. O. Eckert, A. K. Schrey, and R. Preissner, “ProTox-II: a webserver for the prediction of toxicity of chemicals,” Nucleic Acids Res., vol. 46, no. W1, pp. W257–W263, 02 2018, doi: 10.1093/nar/gky318.

[47] J. Machhar, A. Mittal, S. Agrawal, A. M. Pethe, and P. S. Kharkar, “Computational prediction of toxicity of small organic molecules: state-of-the-art,” Phys. Sci. Rev., vol. 4, no. 10, Oct. 2019, doi: 10.1515/psr-2019-0009.

[48] G. Luna, A. V. Dolzhenko, and R. L. Mancera, “Inhibitors of Xanthine Oxidase: Scaffold Diversity and StructureLBased Drug Design,” ChemMedChem, vol. 14, no. 7, pp. 714–743, Apr. 2019, doi: 10.1002/cmdc.201900034.

[49] H. Cao, J. Hall, and R. Hille, “Substrate Orientation and Specificity in Xanthine Oxidase: Crystal Structures of the Enzyme in Complex with Indole-3-acetaldehyde and Guanine,” Biochemistry, vol. 53, no. 3, pp. 533–541, Jan. 2014, doi: 10.1021/bi401465u.

[50] S. Sakkiah, S. Thangapandian, and K. W. Lee, “Ligand-Based Virtual Screening and Molecular Docking Studies to Identify the Critical Chemical Features of Potent Cathepsin D Inhibitors: Pharmacophore-Based Identification of Potent Inhibitor of Cathepsin D,” Chem. Biol. Drug Des., vol. 80, no. 1, pp. 64–79, Jul. 2012, doi: 10.1111/j.1747-0285.2012.01339.x.

[51] D. N. Boobbyer, P. J. Goodford, P. M. McWhinnie, and R. C. Wade, “New hydrogen-bond potentials for use in determining energetically favorable binding sites on molecules of known structure,” J. Med. Chem., vol. 32, no. 5, pp. 1083–1094, May 1989, doi: 10.1021/jm00125a025.

[52] S. Lin, G. Zhang, J. Pan, and D. Gong, “Deciphering the inhibitory mechanism of genistein on xanthine oxidase in vitro,” J. Photochem. Photobiol. B, vol. 153, pp. 463–472, Dec. 2015, doi: 10.1016/j.jphotobiol.2015.10.022.

[53] A. Šmelcerović et al., “Xanthine oxidase inhibitors beyond allopurinol and febuxostat; an overview and selection of potential leads based on in silico calculated physico-chemical properties, predicted pharmacokinetics and toxicity,” Eur. J. Med. Chem., vol. 135, pp. 491–516, Jul. 2017, doi: 10.1016/j.ejmech.2017.04.031.

[54] J. Nomura et al., “Xanthine Oxidase Inhibition by Febuxostat Attenuates Experimental Atherosclerosis in Mice,” Sci. Rep., vol. 4, no. 1, p. 4554, May 2015, doi: 10.1038/srep04554.

[55] A. Rohatgi et al., “HDL Cholesterol Efflux Capacity and Incident Cardiovascular Events,” N. Engl. J. Med., vol. 371, no. 25, pp. 2383–2393, Dec. 2014, doi: 10.1056/NEJMoa1409065.

[56] R. Kumar, Darpan, S. Sharma, and R. Singh, “Xanthine oxidase inhibitors: a patent survey,” Expert Opin. Ther. Pat., vol. 21, no. 7, pp. 1071–1108, Jul. 2011, doi: 10.1517/13543776.2011.577417.

[57] W.-Y. Chung, Y.-J. Jung, Y.-J. Surh, S.-S. Lee, and K.-K. Park, “Antioxidative and antitumor promoting effects of [6]-paradol and its homologs,” Mutat. Res. Toxicol. Environ. Mutagen., vol. 496, no. 1–2, pp. 199–206, Sep. 2001, doi: 10.1016/S1383-5718(01)00221-2.

[58] S. Dugasani, M. R. Pichika, V. D. Nadarajah, M. K. Balijepalli, S. Tandra, and J. N. Korlakunta, “Comparative antioxidant and anti-inflammatory effects of [6]-gingerol, [8]-gingerol, [10]-gingerol and [6]-shogaol,” J. Ethnopharmacol., vol. 127, no. 2, pp. 515–520, Feb. 2010, doi: 10.1016/j.jep.2009.10.004.

[59] S. Prasad and A. K. Tyagi, “Ginger and Its Constituents: Role in Prevention and Treatment of Gastrointestinal Cancer,” Gastroenterology Research and Practice, Mar. 08, 2015. https://www.hindawi.com/journals/grp/2015/142979/ (accessed Jul. 10, 2020).

[60] S. Bais and N. Abrol, “Review on Chemistry and Pharmacological Potential of Amentoflavone,” vol. 6, no. 1, pp. 16–22, 2016, doi: 10.3923/crn.2016.16.22.

[61] J. L. Goldstein and M. S. Brown, “A Century of Cholesterol and Coronaries: From Plaques to Genes to Statins,” Cell, vol. 161, no. 1, pp. 161–172, Mar. 2015, doi: 10.1016/j.cell.2015.01.036.

[62] K. Wang et al., “A novel class of α-glucosidase and HMG-CoA reductase inhibitors from Ganoderma leucocontextum and the anti-diabetic properties of ganomycin I in KK-A y mice,” Eur. J. Med. Chem., vol. 127, pp. 1035–1046, Feb. 2017, doi: 10.1016/j.ejmech.2016.11.015.

[63] D. E. Clark, “In silico prediction of blood–brain barrier permeation,” Drug Discov. Today, vol. 8, no. 20, pp. 927–933, Oct. 2003, doi: 10.1016/S1359-6446(03)02827-7.

[64] M. Muehlbacher, G. M. Spitzer, K. R. Liedl, and J. Kornhuber, “Qualitative prediction of blood–brain barrier permeability on a large and refined dataset,” J. Comput. Aided Mol. Des., vol. 25, no. 12, pp. 1095–1106, Dec. 2011, doi: 10.1007/s10822-011-9478-1.

[65] M. Šrejber et al., “Membrane-attached mammalian cytochromes P450: An overview of the membrane’s effects on structure, drug binding, and interactions with redox partners,” J. Inorg. Biochem., vol. 183, pp. 117–136, Jun. 2018, doi: 10.1016/j.jinorgbio.2018.03.002.

[66] C. C. Ogu and J. L. Maxa, “Drug interactions due to cytochrome P450,” Proc. Bayl. Univ. Med. Cent., vol. 13, no. 4, pp. 421–423, Oct. 2000.

[67] A. Tornio and J. T. Backman, “Cytochrome P450 in Pharmacogenetics: An Update,” in Advances in Pharmacology, vol. 83, Elsevier, 2018, pp. 3–32.

[68] R. Watanabe et al., “Development of an in silico prediction system of human renal excretion and clearance from chemical structure information incorporating fraction unbound in plasma as a descriptor,” Sci. Rep., vol. 9, no. 1, p. 18782, Dec. 2019, doi: 10.1038/s41598-019-55325-1.

[69] M. Bredemeier et al., “Xanthine oxidase inhibitors for prevention of cardiovascular events: a systematic review and meta-analysis of randomized controlled trials,” BMC Cardiovasc. Disord., vol. 18, no. 1, p. 24, Dec. 2018, doi: 10.1186/s12872-018-0757-9.

[70] P. Pacher, A. Nivorozhkin, and C. Szabó, “Therapeutic Effects of Xanthine Oxidase Inhibitors: Renaissance Half a Century after the Discovery of Allopurinol,” Pharmacol. Rev., vol. 58, no. 1, pp. 87–114, Mar. 2006, doi: 10.1124/pr.58.1.6.

[71] P. Pacher, A. Nivorozhkin, and C. Szabó, “Therapeutic Effects of Xanthine Oxidase Inhibitors: Renaissance Half a Century after the Discovery of Allopurinol,” Pharmacol. Rev., vol. 58, no. 1, pp. 87–114, Mar. 2006, doi: 10.1124/pr.58.1.6.

[72] Y. He et al., “Statins and Multiple Noncardiovascular Outcomes: Umbrella Review of Meta-analyses of Observational Studies and Randomized Controlled Trials,” Ann. Intern. Med., vol. 169, no. 8, p. 543, Oct. 2018, doi: 10.7326/M18-0808.

[73] M. Coimbra et al., “Improving solubility and chemical stability of natural compounds for medicinal use by incorporation into liposomes,” Int. J. Pharm., vol. 416, no. 2, pp. 433–442, Sep. 2011, doi: 10.1016/j.ijpharm.2011.01.056.

