## Supplemental file for "*In-Silico* Analysis of Secondary Metabolites that Modulates Enzymes of Cholesterol Target"

**Table 1S.** Details about molecular docking platform

| | |
|---|---|
| Subject | Pharmaceutical Science |
| Specific subject area | Interdisciplinary fields include organic chemistry, biochemistry, and biology. Drug design and discovery from plant sources. |
| Type of data | Tables and Figures |
| How data were acquired | MOE 2009 and GOLD V 4.0.1 |
| Data format | Raw and Analysed |
| Parameters for data collection | Gold Fitness score, energetic values, and interactions of the protein with the ligand. |
| Description of data collection | The protein was collected from the RCSB protein bank. The secondary metabolite structures were obtained from the PubChem online database. The docking was done using GOLD software. |
| Data source location | https://www.rcsb.org/, https://pubchem.ncbi.nlm.nih.gov/ |
| Data accessibility | PDB files of the chosen enzyme targets are publically available at https://www.rcsb.org/ Tables, and Figures of the docking are accessible in the article. |

Value of the Data

- The screening procedure enables the researchers to rapidly identify active natural compounds that can modulate a particular biochemical pathway.
- The screening results help to study the interaction/role of active metabolites in a particular biochemical process at the cellular level and provide preliminary ideas for drug design development
- By using this *in-silico* docking data, novel synthetic analogs with improved bioactivity and minimized side effects can be developed against these targets, and research time can be minimized considerably.
- We select these 16 metabolites because these are abundant in nature and well explored. Among these metabolites, the compounds which show the best affinity for various targets are shortlisted.
- The data is also useful for research scholars who do not have sufficient software and hardware requirements that are not affordable by them.
- Research scholars, researchers in pharmaceutical chemistry, Medicinal Chemistry, Drug Design Industry can benefit from the data.

**Table 2S.** List of Targets showing the PDB ID, resolution and description of the proteins selected for docking with complexed inhibitor

| PDB ID | Resolution (Å) | Description |
|---|---|---|
| 1N5X | 2.8 | Crystal Structure of Xanthine Oxidase from Bovine Milk [1] |
| 1HWK | 2.22 | Crystal structure of human HMG-CoA Reductase with Atorvastatin [2] |

**Table 3S.** Active Site residues of HMG-CoA Reductase and Xanthine Oxidase

| **PDB ID** | **Name of the Organism** | **Active Site Residues** |
|---|---|---|
| 1HWK | Homo sapiens | Arg B568,Ala B856, Ser B565, Leu B853, Leu B562, Ser B852, Gly B560, Asn B755, Lys A692, Ser A661, Cys B561, Glu B559, His B752, Asp A690, Ala B751, Asn A686, Arg A690, Ala B751, Asn A686, Arg A590, Ser A684, Lys B735,Lys A691, Leu B857, Val A683 |
| 1N5X | Bos taurus | Ala 1079, Phe 1009, Arg 880, Ser 1008, Thr 1010, Ser 876, Phe 914, Val 1011, Phe 649, Leu 1014, Leu 648, Pro 1076, Lys 771, Phe 1013, Asn 768, Leu 873, Glu 802,Phe 798, Gln 767, Asp 872, Met 1038 |

**Table 4S.** Molecular Properties of standard compounds and selected secondary metabolites

| Compound Name | Vdw | Elec | Weight | logP | TPSA | Donar | Acceptor |
|---|---|---|---|---|---|---|---|
| Allopurinol | 10.54 | -21.031 | 136.11 | -0.187 | 70.14 | 2 | 3 |
| Amentoflavone **(1)** | 88.370 | -12.116 | 538.464 | 4.820 | 173.980 | 6 | 8 |
| Atorvastatin | 78.69 | -40.826 | 557.64 | 5.245 | 114.62 | 3 | 5 |
| Febuxostat | 33.8 | -8.336 | 315.37 | 2.389 | 86.04 | 0 | 4 |
| Ganoderic acid DM **(15)** | 72.216 | -11.591 | 468.678 | 6.931 | 71.440 | 1 | 4 |
| Ganoderic acid η **(16)** | 81.76 | -15.528 | 531.66 | 1.479 | 155.19 | 4 | 8 |
| Ganoleucoin K **(11)** | 85.021 | --20.835 | 670.752 | 0.730 | 215.30 | 2 | 11 |
| Ganoleucoin T **(14)** | 84.62 | -2.034 | 597.76 | 3.56 | 148.87 | 2 | 8 |
| Ganoleucoin Y **(13)** | 98.04 | -31.164 | 672.76 | 0.521 | 218.46 | 3 | 11 |
| Ganoleucoin Z **(12)** | 101 | -29.294 | 670.75 | 0.73 | 215.3 | 2 | 11 |
| Ganomycin I **(9)** | 32.44 | -7.947 | 342.43 | 5.191 | 66.76 | 2 | 3 |
| Hydroxychavicol **(7)** | 19.804 | -2.997 | 150.177 | 1.826 | 40.46 | 2 | 2 |
| Isoquercitrin **(5)** | 76.126 | -24.94 | 463.37 | -0.292 | 209.43 | 7 | 11 |
| Lovastatin | 42.59 | -11.604 | 404.54 | 4.196 | 72.83 | 1 | 3 |
| Neotaiwanensol B **(6)** | 46.12 | -3.304 | 298.33 | 3.555 | 80.92 | 4 | 4 |
| Pravastatin | 52.7 | -39.373 | 423.52 | 1.106 | 127.12 | 3 | 6 |
| Probenecid | 28.09 | 1.256 | 284.35 | 0.861 | 77.51 | 0 | 4 |
| Riparsaponin **(8)** | 87.136 | 21.061 | 620.868 | 3.791 | 139.840 | 6 | 8 |

| | | | | | | | |
|---|---|---|---|---|---|---|---|
| Selgin **(4)** | 55.268 | -31.384 | 316.265 | 2.134 | 116.450 | 4 | 6 |
| Simvastatin | 47.09 | -11.683 | 418.57 | 4.586 | 72.83 | 1 | 3 |
| Topiroxostat | 33.07 | 3.108 | 248.24 | 1.8 | 91.14 | 1 | 5 |
| n-octadecanyl-O-**α**-D-glucopyranosyl(6'→1")-O-**α**-D-glucopyranoside **(10)** | 55.396 | 10.940 | 594.783 | 1.889 | 178.530 | 7 | 11 |
| 6- Gingerol **(2)** | 35.09 | -10.246 | 294.39 | 3.234 | 66.76 | 2 | 4 |
| 6- Paradol **(3)** | 33.14 | -6.873 | 278.39 | 4.263 | 46.53 | 1 | 3 |

Where: Vdw = Van der waals energy, Elect = Electrostatic energy, logP =Partition coefficient, TPSA =Total polar surface area

**Table 5S.** Inhibition of HMG-CoA reductase by secondary metabolites

| **Natural Source** | **Secondary metabolite** | **$IC_{50}$ value** | **Reference** |
|---|---|---|---|
| *Ficus virens*<br>Bark | n-octadecanyl-O-α-D-glucopyranosyl(6'→1")-O-α-D-glucopyranoside **(10)*** | 0.164 µM | [3] |
| *Ganoderma leucocontextum* | Ganoleucoin Z **(12)***<br>Ganoleucoin Y **(13)***<br>Ganoleucoin T **(14)*** | 8.68 ± 0.96 µM<br>9.72 ± 0.91µM<br>10.3 ± 1.78 µM | [4] |
| *Ganoderma leucocontextum*<br>Fruit | Ganoderic acid DM **(15)**<br>Ganoleucoin K **(11)**<br>Ganoderiol J | 9.5 ± 1.5 µM<br>10.7 ± 2.9 µM<br>12.6 ± 2.7 µM | [5] |
| *Ganoderma leucocontextum*<br>Fruit | Ganomycin I **(9)**<br>Ganomycin B<br>Ganomycin C<br>Fornicin B | 12.3 ± 1.7 µM<br>29.3 ± 2.5 µM<br>45.2 ± 7.1 µM<br>56.9 ± 12.1 µM | [6] |
| *Ganoderma lucidum*<br>Fruit | Ganomycin I<br>Ganoderenic acid K<br>Ganoderic acid η **(16)**<br>Ganomycin B | 14.3 ± 1.5 µM<br>16.5 ± 2.4 µM<br>29.8 ± 1.5 µM<br>30.3 ± 1.5 µM | [7] |
| *Ganoderma Lucidum* | 15-hydroxy-ganoderic acid S<br>7-oxo-ganoderic acid Z | 21.7 µM<br>22.3 µM | [8] |
| *Vitis vinifera*<br>Stem bark | Vitisin B<br>Vitisin A<br>γ-Viniferin<br>Ampelopcin-A | 23.9 ± 5.0 µM<br>42 ± 3.1 µM<br>232.6 ± 30.9 µM<br>294 ± 9.5 µM | [9] |
| *Rosa damascena* | Roxyloside<br>Quercetin gentiobioside<br>Afzelin<br>Isoquercitrin | 47.1 µM<br>50.6 µM<br>80.1 µM<br>80.6 µM | [10] |

Note: Some $IC_{50}$ values are adjusted in terms of molarity to make the comparison convenient; *Already docked molecules
Current medication: Atorvastatin, Simvastatin, Pravastatin, Lovastatin

**Table 6S.** Inhibition of Xanthine Oxidase by secondary metabolites

| Natural Source | Secondary Metabolites | $IC_{50}$ value | References |
|---|---|---|---|
| *Homonoia riparia* Lour | Riparsaponin **(8)** | 0.011 µM | [11] |
| *Semecarpus anacardium* | Amentoflavone **(1)** | 0.092 µM | [12] |
| *Chrysanthemum sinense* | Diosmetin* | 0.13 µM | [13] |
| | Acacetin* | 0.16 µM | |
| | Chrysoeriol | 0.19 µM | |
| | Eupafolin | 0.20 µM | |
| | Selgin **(4)** | 0.22 µM | |
| | Apigenin | 0.36 µM | |
| | Jaceidin | 1.15 µM | |
| | Luteolin* | 1.24 µM | |
| | 4,5-O-dicaffeoylquinic acid methyl ester | 2.31 µM | |
| *Piper nudibaccatum* | Neotaiwanesol B **(6)** | 0.28 µM | [14] |
| | Hydroxychavicol **(7)*** | 0.38µM | |
| Flavones (class) | Isorhamnetin* | 0.4 µM | [15] |
| | Kaempferol* | 0.67 µM | |
| | Myricetin* | 1.27 µM | |
| | Rutin* | 46.8 µM | |
| | Genistein* | 83 µM | |
| *Perilla frutescens* | Apigenin* | 0.44 µM | [16] |
| | Methyl rosamarinate | 26.59 µM | |
| | Vinyl caffeate | 31.26 µM | |
| | Rosamarinic acid* | 91.72 µM | |
| | Caffeic acid* | 121.22 µM | |
| Dihydrochalcone (Class) | Phloretin* | 0.66 µM | [15] |
| Coffee beans | Pyrogallol* | 0.73 µM | [17] |

| | | | |
|---|---|---|---|
| *Stauntonia brachyanthera* | Isoquercitrin **(5)** | 1.6 μM | [18] |
| | 3β,20α,24-trihydroxy-29-norolean-12-en-28-oic-acid-24-O-β-L-fucopyranosyl- (1→ 2)-6-O-acetyl-β-D-glucopyranoside | 5.22 μM | |
| *Blumea balsamifera* | Luteolin* | 2.38 μM | [19] |
| | Quercetin* | 2.92 μM | |
| | Tamarixetin* | 3.16 μM | |
| | 5,7,3',5'-Tetrahydroxy flavone | 32.14 μM | |
| | Rhamnetin* | 36.09 μM | |
| | Blumeatin | 53.21 μM | |
| | Dihydroquercetin-4'-methyl ether | 58.46 μM | |
| | Luteolin-7-methyl ether | 42.19 μM | |
| *Lagerstroemi speciosa* | Valoneic acid | 2.5 μM | [20] |
| | Ellagic acid * | 71.5 μM | |
| *Toona sinensis* | 1,2,3,4,6-Penta-O-galloyl-*β*-D-glucopyranose | 2.8 μM | [21] |
| *Rabdosia japonica* Hara | 3,4-Dihydroxy phenyl acetic acid | 3.5 ± 0.5 μM | [22] |
| *Centaurea virgata* Lam. | Hispidulin | 4.88 μM | [23] |
| *Amentotaxus formosana* | (+) Sugiol | 6.8 ± 0.4 μM | [24] |
| *Zea mays* L. | Ferulic acid* | 8.2 ± 0.3 μM | [25] |
| *Momordica charantia* | Esculetin* | 8.2 μM | [26] |
| | Taiwacin A | 24.3 ± 3.4 μM | |
| *Salviae miltiorrhizae* | Lithospermic acid* | 9.65 μM | [27] |
| *Zingiber officinale* | 6-Gingerol **(2)*** | 10.5 μM | [28] |
| | 6-Paradol **(3)** | 12.4 μM | |
| | 6-Shogaol* | 15.2 μM | |
| *Herpetospermum penduculosum* | Cucurbitacin E | 10.16 ± 0.21 μM | [29] |
| | Neocucurbitacin D | 15.27 ± 0.29 μM | |
| | Cucurbitacin B* | 18.41 ± 0.34 μM | |
| *Veratrum taliense* | Veraphenol | 11 μM | [30] |
| | Piceid | 66.1 μM | |

| | | | |
|---|---|---|---|
| | Isorhapontin | 70 μM | |
| | Mulberroside E | 78.4 μM | |
| | Resveratol* | 96.7 μM | |
| *Pueraria lobata* | Genistein* | 11.18 μM | [31] |
| | Diadzin | 12.75 μM | |
| | Daidzein* | 57.03 μM | |
| | Puerarin | 73.96 μM | |
| *Pistacia integerrima* | Quercetin-3-O-**β**-D-glucopyranoside | 12.389 μM | [32] |
| | Kaempferol-3-O-**β**-D-glucopyranoside | 26.199 μM | |
| *Carallia brachiate* | Carallidin | 12.9 μM | [33] |
| *Rhodiola crenulata* | 4'-Hydroxy acetophenone | 15.62 ± 1.19 μM | [34] |
| | Epicatechin-(4β,8)-epicatechin gallate | 24.24 ±1.8 μM | |
| *Citrus aurantium* | Hesperetin* | 16.48 μM | [35] |
| | Nobiletin | 107.51 μM | |
| *Piper betel* | 4-Allyl-1,3-hydroxybenzene (hydroxychavicol) | 16.7 μM | [36] |
| Radix Salviae | Danshenxinkun B | 17.45 ± 2.1μM | [27] |
| *Paulownia catalpifolia* | Paucatalinone N | 20.3 μM | [37] |
| | Paucatalinone L | 29.6 μM | |
| Anthocyanidin (Class) | Pelargonidin | 21.9 μM | [15] |
| | Peonidin | 26.0 μM | |
| | Cyanidin | 27.8 μM | |
| | Apigenidin | 29.1 μM | |
| | Delphinidin | 52.4 μM | |
| *Palhinhaea ceruna* | Apigenin-4'-O-(2' '-O-p-coumaroyl)-β-d-glucopyranoside | 23.95 μM | [38] |
| *Centaurium erythraea* | Caulerpenyne | 26.92 μM | [39] |
| | Sinapic acid | 147.4 μM | |
| *Artocarpus communis* | Artonol A | 43.3 ± 8.1 μM | [40] |
| | Cyclogera communin | 73.3 ± 19.1 μM | |

| | | | |
|---|---|---|---|
| *Morus alba* L. | Morin | 44 µM | [41] |
| *Sinofranchetia chinensis* | Liquiritigenin | 49.3 µM | [42] |
| | Isoliquiritigenin | 55.8 µM | |
| *Garcinia subelliptica* | Garcinielliptones | 53.8 ± 11.5 µM | [43] |
| *Scutellaria baicalensis* | Baicalin* | 55.58 µM | [44] |
| *Cinnamomum osmophloeum* | Cinnamaldehyde* | 63.55 µM | [45] |
| *Coumarin derivatives* | 3-Hydroxy coumarin | 131 µM | [46] |
| | 7-Hydroxy-7-methoxy coumarin | 138 µM | |
| | 4- Methyl esculetin* | 246 µM | |
| - | p-Coumaric acid* | 144.9 µM | [47] |
| - | Limonene* | 308.96 µM | |
| - | β-Caryophyllene* | 319.48 µM | |

Note: Some $IC_{50}$ values are adjusted in terms of molarity to make the comparison convenient; *Already docked molecules
Current medication: Allopurinol, Febuxostat, Pegloticase, Probenecid

**Table 7S.** Prediction of toxicity of secondary metabolites inhibiting metabolic enzymes using ProTox-II

| **Compound** | **$LD_{50}$ mg/Kg** | **Toxicity class** | **Active target** | **Probability** |
|---|---|---|---|---|
| Alisol A acetate | 5000 | 5 | Immunotoxicity<br>Androgen receptor | 0.82<br>0.86 |
| Alisol B acetate | 15070 | 6 | Immunotoxicity<br>Androgen receptor | 0.96<br>0.81 |
| Allopurinol | 78 | 3 | Hepatotoxicity | 0.73 |
| Amentoflavone **(1)** | - | 4 | - | - |
| Ampleopcin-A | 2000 | 4 | Aryl hydrocarbon receptor<br>Estrogen receptor alpha<br>Mitochondrial membrane<br>potential | 0.70<br><br>0.70<br>0.84 |
| Atorvastatin (Commercial drugs for HMG CoA- reductase) | 5000 | 5 | Hepatotoxicity<br>Aromatase | 0.74<br>0.98 |
| Febuxostat (Commercial drugs for XO) | 8000 | 6 | Hepatotoxicity<br>Aryl hydrocarbon Receptor (AhR)<br>Androgen Receptor (AR)<br>ATPase family AAA domain-containing protein 5 (ATAD5) | 0.83<br>1<br>1<br>1 |
| Fornicin B | 1000 | 4 | Immunotoxicity | 0.7 |
| Ganoderenic acid K | 3000 | 5 | Immunotoxicity<br>Androgen receptor | 0.99<br>0.78 |
| Ganoderic acid DM **(15)** | 1185 | 4 | - | - |
| Ganoderic acid η **(16)** | 9000 | 6 | Immunotoxicity<br>Androgen receptor | 0.99<br>0.75 |
| Ganoleucoin K **(11)** | 200 | 3 | Androgen receptor<br>Androgen receptor ligand binding domain | 0.97<br>0.97 |
| Ganoleucoin T **(14)** | 9000 | 6 | Immunotoxicity<br>Androgen receptor<br>Androgen receptor ligand binding domain | 0.93<br>0.90<br>0.96 |
| Ganoleucoin Y **(13)** | 79 | 3 | Androgen receptor<br>Androgen receptor ligand binding domain | 0.97<br>0.98 |

| | | | | |
|---|---|---|---|---|
| Ganoleucoin Z **(12)** | 79 | 3 | Androgen receptor<br>Androgen receptor ligand binding domain | 0.97<br>0.98 |
| Ganomycin I **(9)** | 2000 | 4 | - | - |
| Ganomycin B | 370 | 4 | - | - |
| Ganomycin C | 400 | 4 | Immunotoxicity | 0.93 |
| Hydroxychavicol **(7)** | 1930 | 4 | - | - |
| Isoliquiritigenin | 1048 | 4 | Immunotoxicity<br>Estrogen Receptor Alpha (ER)<br>Estrogen Receptor Ligand Binding Domain (ER-LBD)<br>Mitochondrial Membrane Potential (MMP)<br>Phosphoprotein (Tumor Suppressor) p53<br>ATPase family AAA domain-containing protein 5 (ATAD5) | 0.79<br>1<br>1<br>0.99<br>1<br>1 |
| Isoquercitrin **(5)** | 5000 | 5 | - | - |
| Kaempferol-3-O-beta-D-glucopyranoside | 5000 | 5 | - | - |
| Liquiritigenin | 2000 | 4 | Estrogen Receptor Alpha (ER)<br>Domain (ER-LBD)<br>Mitochondrial Membrane Potential (MMP)<br>ATPase family AAA domain-containing protein 5 (ATAD5) | 0.96<br>0.81<br>0.76<br>0.70 |
| Lunatinin | 1000 | 4 | Mitochondrial membrane potential | 0.74 |
| Lycopene | 5000 | 5 | Androgen receptor ligand binding<br>Estrogen receptor<br>Alpha Estrogen receptor ligand binding domain | 1.0<br>1.0<br>1.0 |
| Neotaiwanensol B **(6)** | 1500 | 4 | Mitochondrial Membrane Potential (MMP) | 0.83 |
| n-octadecanyl-O-α-D-glucopyranosyl(6’ →1”)-O-α-D-glucopyranoside **(10)** | 2000 | 4 | - | - |
| Pravastatin Commercial drugs for HMG CoA- reductase) | 8939 | 6 | Immunotoxicity | 0.97 |
| Quercetin-3-O-beta-D-glucopyranoside | 5000 | 5 | - | - |

| | | | | |
|---|---|---|---|---|
| Riparsaponin **(8)** | 1966 | 5 | - | - |
| Selgin **(4)** | 8357 | 5 | - | - |
| Vitisin A | 1340 | 4 | Immunotoxicity<br>Mitochondrial membrane potential | 0.86<br>0.73 |
| Vitisin B | 1340 | 4 | Immunotoxicity<br>Aryl hydrocarbon receptor<br>Mitochondrial membrane potential | 0.96<br>0.74<br>0.78 |
| 3ß,20a,24-trihydroxy-29-norolean-12-en-28-oic acid 24-O-ß-L-fucopyranosyl- (1→2)-6-O-acetyl-ß-D-glucopyranoside | 8000 | 6 | Immunotoxicity | 0.99 |
| ϒ-viniferin | 1743 | 4 | Immunotoxicity<br>Aryl hydrocarbon receptor<br>Estrogen receptor alpha<br>Mitochondrial membrane potential | 0.75<br>0.75<br>0.73<br>0.91 |
| 6-Gingerol **(2)** | 250 | 3 | Immunotoxicity | 0.96 |
| 6- Paradol **(3)** | 258 | 5 | - | - |
| 6- Shogaol | 687 | 4 | Immunotoxicity | 0.86 |

**Table 8S (a):** ADMET properties of XO inhibitors by pKCSM server

| | Parameters | 1` | 2 | 3 | 4 | 5 | 6 | 7 | 8 | Febuxostat | Probenecid | Allopurinol |
|---|---|---|---|---|---|---|---|---|---|---|---|---|
| Absorption | Water solubility (log mol/L) | -2.892 | -3.232 | -4.186 | -3.38 | -4.07 | -3.528 | -2.898 | -1.252 | -3.019 | -2.458 | -2.378 |
| | Caco2 permeability (log Papp 10-6 cm/s) | -0.183 | 0.776 | 1.537 | 0.121 | 0.545 | 1.157 | -0.779 | 1.689 | 1.031 | 0.997 | 0.503 |
| | Intestinal absorption (% absorbed) | 80.183 | 90.369 | 91.07 | 74.744 | 68.288 | 89.771 | 44.424 | 90.549 | 93.929 | 97.602 | 94.177 |
| | Skin permeability (log Kp) | -2.735 | -2.911 | -2.592 | -2.735 | -2.719 | -2.735 | -2.735 | -2.473 | -2.734 | -2.73 | -2.737 |
| Distribution | VDss (Human, log L/Kg) | -1.177 | 0.08 | 0.244 | 0.454 | -1.121 | -0.256 | 1.179 | 0.249 | -1.209 | -1.466 | 0.056 |
| | BBB Permeability (logBB) | -1.836 | -0.511 | -0.184 | -1.164 | -1.058 | -0.857 | -1.794 | 0.149 | -0.629 | -0.56 | -1.393 |
| | CNS Permeability (log PS) | -3.499 | -2.735 | -1.572 | -3.193 | -3.091 | -2.293 | -4.663 | -2.022 | -2.146 | -2.935 | -4.57 |
| Metabolism | CYP1A2 | NO | NO | YES | YES | NO | YES | NO | YES | NO | NO | NO |
| | CYP2C19 | NO | YES | YES | NO | NO | YES | NO | NO | NO | NO | NO |
| | CYP2C9 | NO | NO | NO | NO | NO | YES | NO | NO | NO | NO | NO |
| | CYP2D6 | NO | NO | NO | NO | NO | NO | NO | NO | NO | NO | NO |
| | CYP3A4 | NO | NO | NO | NO | NO | NO | NO | NO | NO | NO | NO |
| Excretion | Renal OCT2 substrate clearance | NO | NO | NO | NO | NO | NO | NO | NO | NO | NO | NO |
| | Total Clearance (logml/min/kg) | 0.502 | 1.369 | 1.428 | 0.559 | 0.419 | 0.151 | 0.206 | 0.296 | 0.313 | 1.058 | 0.57 |
| Toxicity | Ames Toxicity | NO | YES | NO | NO | NO | NO | NO | YES | NO | NO | NO |
| | Hepatotoxicity | NO | NO | NO | NO | NO | NO | NO | NO | NO | NO | NO |
| | Rat Oral Toxicity (LD50) | 2.549 | 2.06 | 2.249 | 2.41 | 4.352 | 2.436 | 2.545 | 2.165 | 2.649 | 2.438 | 2.3 |
| | Toxicity class | 4 | 3 | 5 | 5 | 5 | 4 | 4 | 5 | 6 | 4 | 6 |

**Table 8S (b):** ADMET properties of HMGR inhibitors by pKCSM server

| | Parameters | 9 | 10 | 11 | 12 | 13 | 14 | 15 | 16 | Atorva statin | Simvast atin | Pravast atin | Lovastatin |
|---|---|---|---|---|---|---|---|---|---|---|---|---|---|
| Absorption | Water solubility (log mol/L) | -4.529 | -2.28 | -3.165 | -3.17 | -3.168 | -3.934 | -4.612 | -4.063 | -3.962 | -4.987 | -2.903 | -4.73 |
| | Caco2 permeability (log Papp 10-6 cm/s) | 0.691 | -0.914 | -0.248 | -0.28 | -0.263 | -0.136 | 0.664 | -0.164 | 0.805 | 0.977 | 0.683 | 0.985 |
| | Intestinal absorption (% absorbed) | 89.814 | 15.261 | 25.85 | 21.02 1 | 14.907 | 59.842 | 97.205 | 46.187 | 93.173 | 95.313 | 41.007 | 95.702 |
| | Skin permeability (log Kp) | -2.881 | -2.735 | -2.735 | -2.735 | -2.735 | -2.735 | -2.733 | -2.738 | -2.735 | -3.347 | -2.83 | -3.364 |
| Distributi | VDss (Human, log L/Kg) | 0.268 | -0.612 | -0.514 | -0.574 | -0.818 | -1.065 | -0.819 | -0.9 | -1.346 | 0.197 | -0.508 | 0.175 |
| | BBB Permeability (logBB) | -0.393 | -1.735 | -1.706 | -1.844 | -1.531 | -0.96 | -0.059 | -1.055 | -1.264 | -0.37 | -1.273 | -0.375 |
| | CNS Permeability (log PS) | -1.979 | -4.863 | -3.221 | -3.337 | -3.433 | -2.569 | -1.425 | -3.103 | -2.379 | -2.812 | -3.403 | -2.832 |
| Metabolism | CYP1A2 | YES | NO | NO | NO | NO | NO | NO | NO | NO | NO | NO | NO |
| | CYP2C19 | YES | NO | NO | NO | NO | NO | NO | NO | YES | NO | NO | NO |
| | CYP2C9 | YES | NO | NO | NO | NO | NO | NO | NO | YES | NO | NO | NO |
| | CYP2D6 | NO | NO | NO | NO | NO | NO | NO | NO | NO | NO | NO | NO |
| | CYP3A4 | NO | NO | NO | NO | NO | NO | NO | NO | YES | YES | NO | YES |
| Excretion | Renal OCT2 substrate clearance | NO | NO | NO | NO | NO | NO | NO | NO | NO | NO | NO | NO |
| | Total Clearance (logml/min/kg) | 1.084 | 2.04 | 0.073 | 0.066 | 0.129 | 0.371 | 0.277 | 0.435 | 0.696 | 0.827 | 1.246 | 0.928 |
| Toxicity | Ames Toxicity | NO | NO | NO | NO | NO | NO | NO | NO | NO | NO | NO | NO |
| | Hepatotoxicity | NO | NO | NO | NO | NO | YES | NO | NO | YES | NO | NO | NO |
| | Rat Oral Toxicity (LD50) | 2.047 | 2.295 | 2.912 | 2.908 | 2.904 | 3.382 | 2.378 | 3.044 | 3.233 | 2.14 | 3.266 | 2.119 |
| | Toxicity class | 4 | 4 | 3 | 3 | 3 | 6 | 4 | 6 | 5 | 4 | 6 | 4 |

**Value Range : LogS (Solubility)**: Optimal (higher than -4log mol/L) , **Papp (Caco-2 Permeability):** Optimal (higher than -5.15 Log unit or -4.70 or -4.80) , **HIA (Human Intestinal Absorption)**: >30% Perfectly absorbed , **VD (Volume Distribution)**: Optimal (0.04-20 L/Kg) , **BBB (Blood Brain Carrier):** ( BB ratio >=0.1: BBB+ ; BB ratio <0.1: BBB-) , **Total Clearance:** >15 ml/min/kg: High; 5 ml/min/kg< CL< 15ml/min/kg: Moderate; <5 ml/min/kg: Low, **LD50( LD50 of acute toxicity):** High-toxicity: (1-50 mg/kg); Modarate-toxicity: (51-500 mg/kg); Low-Toxicity:

(501-5000 mg/kg) , **ProTox-II toxicity class**: Class 1, 2: Fatal; Class 3: Toxic; Class 4, 5: Harmful; Class 6: Non-toxic

**Table 9S.** Crude natural product extracts showing the inhibitory activity against HMGR and XO

| Enzyme | Natural Source | Crude extract | Parts used | $IC_{50}$ value | References |
|---|---|---|---|---|---|
| **HMG-CoA reductase** | *Syzygium polyanthum* | Ethanolic extract | Leaves | Percolation: 49.5 µg/mL<br>Soxhlet: 15.50 µg/mL<br>Simvastatin: 0.00238µg/mL | [48] |
| | *Citrus paradissi* | Phenolic extract | Peels | 114.99 µg/L | [49] |
| | *Basella alba* | Methanolic extract | Leaves | 74.1% inhibition | [50] |
| | *Amaranthus viridis* | Leaf extract | Leaves | 72% inhibition | [51] |
| | *Pithecellobium ellipticum* | Ethanolic extract | Fruits | 80.9% | [52] |
| | *Gnetum gnemon* | Dichloromethane extract | Seeds | (64.78% at 100ppm)<br>0.40 µg/mL | [53] |
| | *Quercus infectoria* | Methanolic extract | Galls | 84% | [54] |
| | *Rosa damascena* | Ethanolic extract | Floret | 70% | |
| | *Myrtus communis* | Methanolic extract | Leaves | 62% | |
| | *Ficus palmata* Forsk | Aqueous extract | Stem bark | 9.1 ± 0.6 µg/mL | [55] |
| | *Guazuma ulmifolia* | Ethanolic extract | Leaves | 69.10 % | [56] |
| | *Guazuma ulmifolia* | Methanolic extract | Leaves | (79.85- 94.42) % | [57] |
| | *Antocephalus macrophyllus* | Methanolic extract | Leaves | (69.96- 91.25) % | |
| | *Melia azedarach* | Methanolic extract | Leaves | (63.12- 96.45) % | |
| **Xanthine oxidase** | *Strychnos nux-vomica L.* | Methanolic extract | Leaves | 6.80 µg/mL<br>Allopurinol: 6.75 µg/mL | [58] |
| | *Tribulus arabicus* | Ethanolic extract | Aerial parts | 20.4 µg/mL<br>Allopurinol: 6.5 µg/mL | [59] |
| | *Artemisia vulgaris* | Methanolic extract | Aerial parts | 14.7 µg/mL<br>Allopurinol: 0.28 µg/mL | [60] |
| | *Pistacia lentiscus* | Aqueous extract from hexane partitions | Leaves | 72.74 ± 2.63% at 100 µg/mL | [61] |

| | | | | |
|---|---|---|---|---|
| *Salvia spinosa* L. | Methanol extract | - | 53.7 µg/mL | [62] |
| *Anthemis palestina* Boiss. | Methanol extract | - | 168.0 µg/mL | |
| *Chrysanthemum coronarium* L. | Methanol extract | - | 199.5 µg/mL | |
| *Ginkgo biloba* L. | Methanol extract | - | 595.8 µg/ml | |
| *Achillea biebersteinii* Afansiev | Methanol extract | - | 360.0 µg/mL | |
| *Rosmarinus officinalis* L. | Methanol extract | - | 650.0 µg/mL | |
| *Cinnamomum cassia* | methanol extract | Twigs | 18 µg/mL | [63] |
| *Chrysanthemum indicum* | methanol extract | Flowers | 22 µg/mL | |
| *Lycopus europaeus* | methanol extract | Leaves | 26 µg/mL | |
| *Polygonum cuspidatum* | Aqueous extract | Rhizome | 38 µg/mL | |
| *Populus nigra* | Methylene chloride–methanolic extracts | Leaves | 8.3 µg/mL | [64] |
| *Betula pendula* | Methylene chloride–methanolic extracts | Leaves | 25.9 µg/mL | |
| *Alocasia longiloba* | Ethanolic extract | Petioles | 42.71 µg/mL | [65] |
| | | Fruits | 51.32 µg/mL | |

**Table 10S.** Gold Fitness score and Protein-Ligand Interactions of Protein ID: 1HWK, HMG-CoA Reductase Inhibition, and Protein ID: 1N5X Xanthine Oxidase Inhibition. The Gold Fitness score, interacting residues, type of interaction, bond length between residues, and ligands are shown.

| S.No. | Compounds | Gold Score | H-Bond Interaction Residues | Bond Length | Other Interacting Residues |
|---|---|---|---|---|---|
| 1 | Atorvastatin (HMG-CoA) | 73.24 | Ser B565 | 2.7 | Pi-Pi interaction with Arg A590 |
| | | | Lys A691 | 2.8 | |
| | | | Asp A690 | 1.6 | |
| | | | Arg A590 | 3 | |
| | | | Ser A684 | 1.9 | |
| | | | Lys B735 | 2.4 | |
| 2 | Febuxostat (XO) | 64.53 | Arg 880 | 2.3 | - |
| | | | Thr 1010 | 2 | |
| | | | Phe 914 | 2 | |
| | | | Asn 768 | 3.5 | |
| 3 | Topiroxostat (XO) | 61.46 | Glu 1261 | 1.90,2.30 | Pi-Pi interaction with Phe 914 |
| | | | Phe 914 | - | |
| 4 | Probenecid (XO) | 57.75 | Met 1038 | 2.5 | - |
| 5 | Simvastatin (HMG-CoA) | 56.81 | Arg A590 | 2.6 | - |
| | | | Ser A684 | 3 | |
| 6 | Pravastatin (HMG-CoA) | 54.83 | Lys A691 | 2.5 | - |
| | | | Glu B559 | 1.9 | |
| | | | Asn B755 | 2.5 | |
| | | | Asp A690 | 1.8 | |
| | | | Arg A590 | 3.10, 3.10 | |
| | | | Lys B735 | 2.9 | |
| | | | Ser A684 | 1.8 | |
| 7 | Allopurinol (XO) | 46.16 | Phe 914 | - | Pi-Pi interaction with Phe 914 |
| | | | Glu 802 | 2.1 | |

| | | | | | |
|---|---|---|---|---|---|
| | | | Thr 1010 | 2.9 | |
| 8 | Lovastatin (HMG-CoA) | 41.36 | Arg A590 | 3 | - |
| | | | Ser A664 | 3.6 | |
| | | | Lys A692 | 2.8 | |

**Note:** Atorvastatin, Simvastatin, Pravastatin,and Lovastatin are commercial HMG-CoA reductase inhibitors while Allopurinol, Febuxostat, and Probenecid  are commercial XO inhibitors

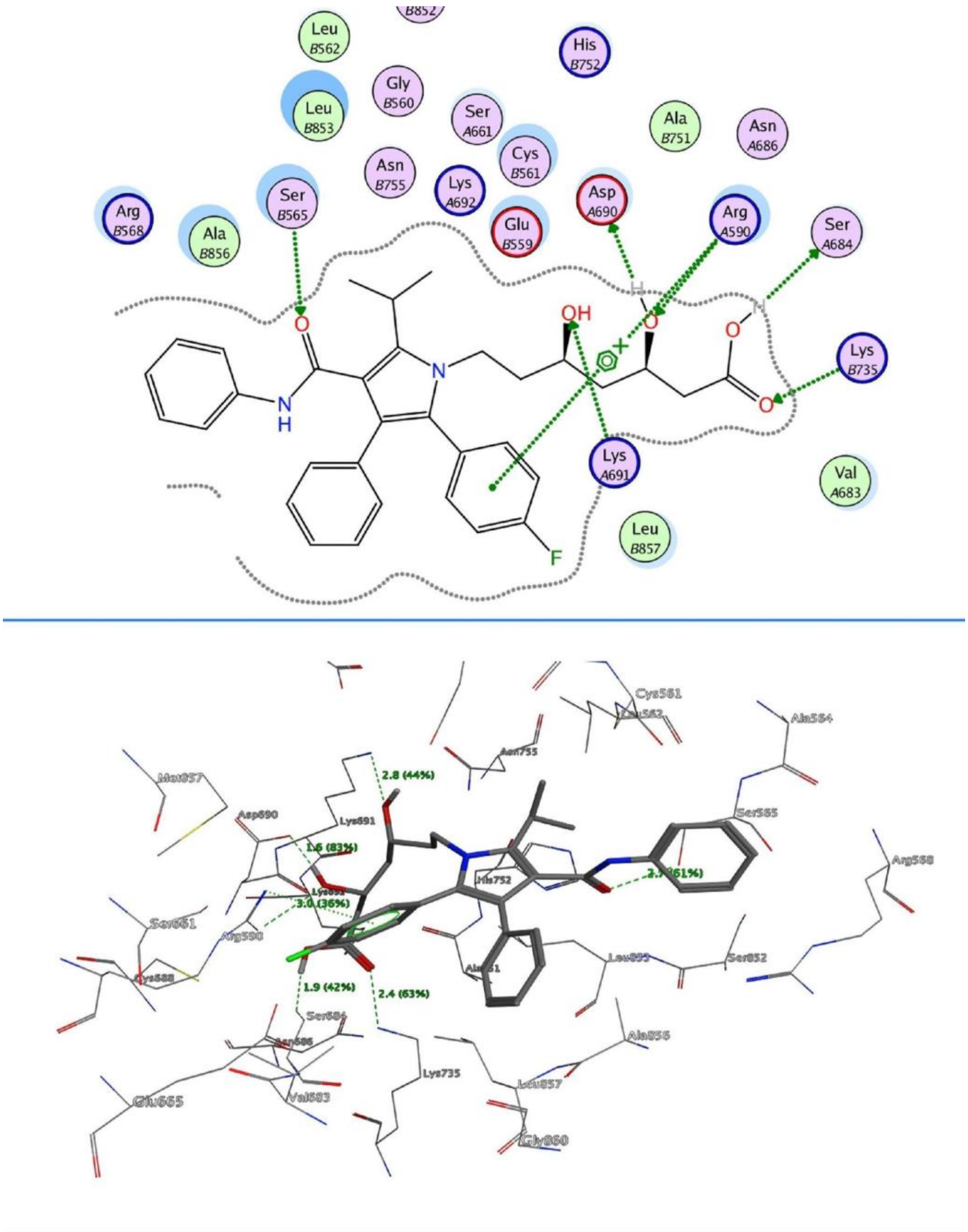


**Figure 1S**. 2D (upper) and 3D (lower) interactions of HMG-CoA Reductase (PDB ID: 1HWK) with atorvastatin (Fitness score of 73.24).

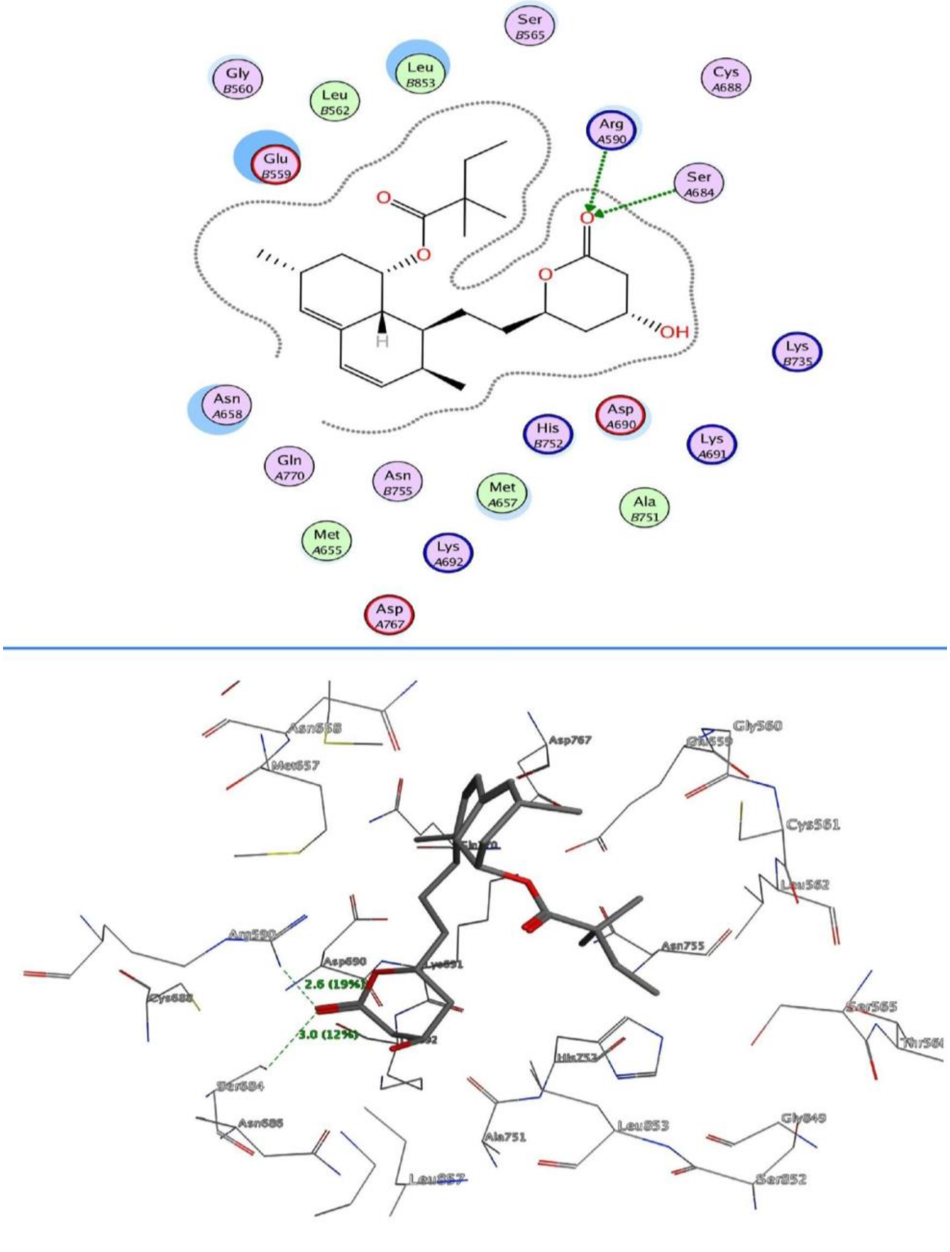


**Figure 2S.** 2D (upper) and 3D (lower) interactions of HMG-CoA Reductase with simvastatin (Fitness score of 56.81).

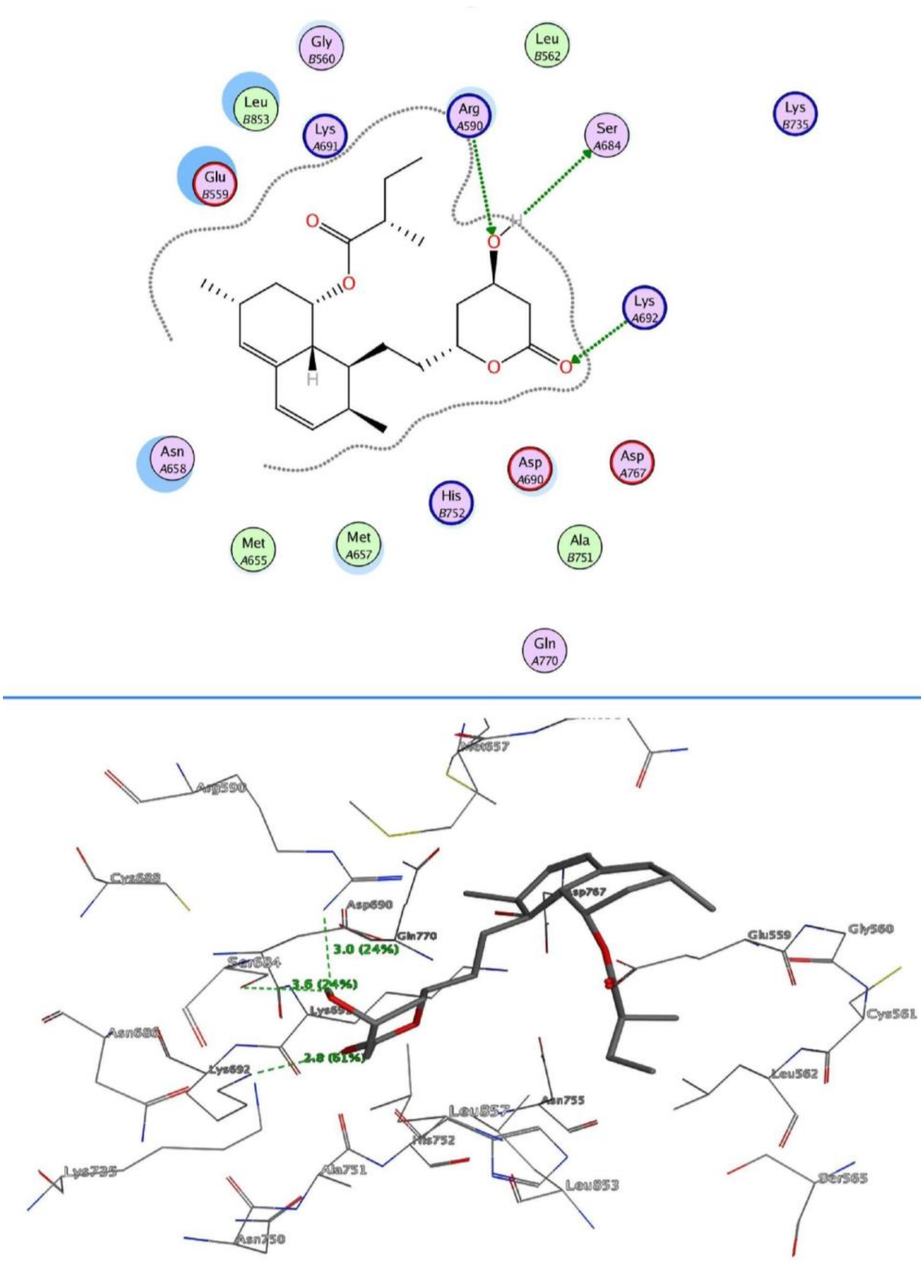


**Figure 3S.** 2D (upper) and 3D (lower) interactions of HMG-CoA Reductase with lovastatin (Fitness score of 41.36).

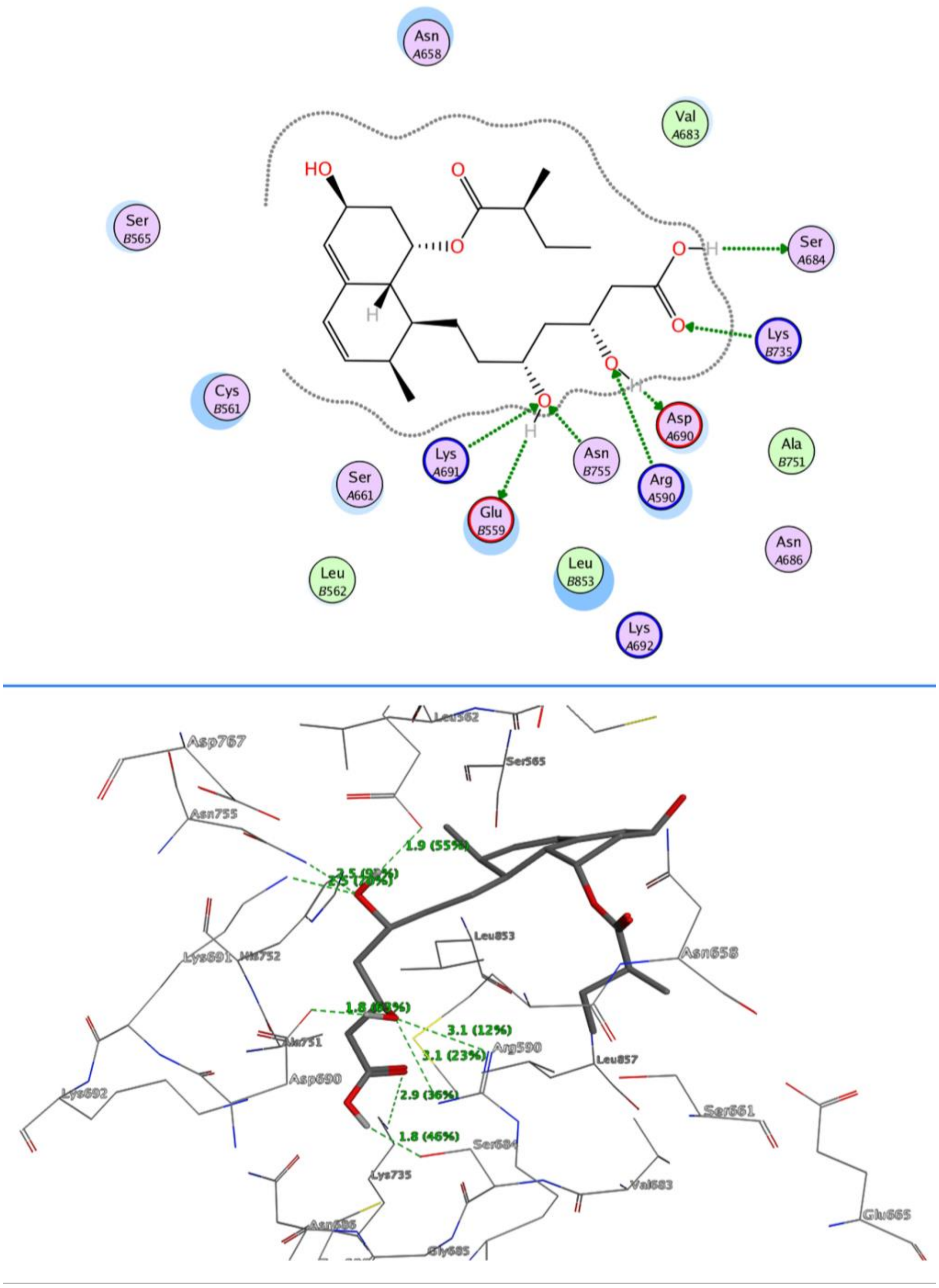


**Figure 4S.** 2D (upper) and 3D (lower) interactions of HMG-CoA Reductase with pravastatin (Fitness score of 54.83).

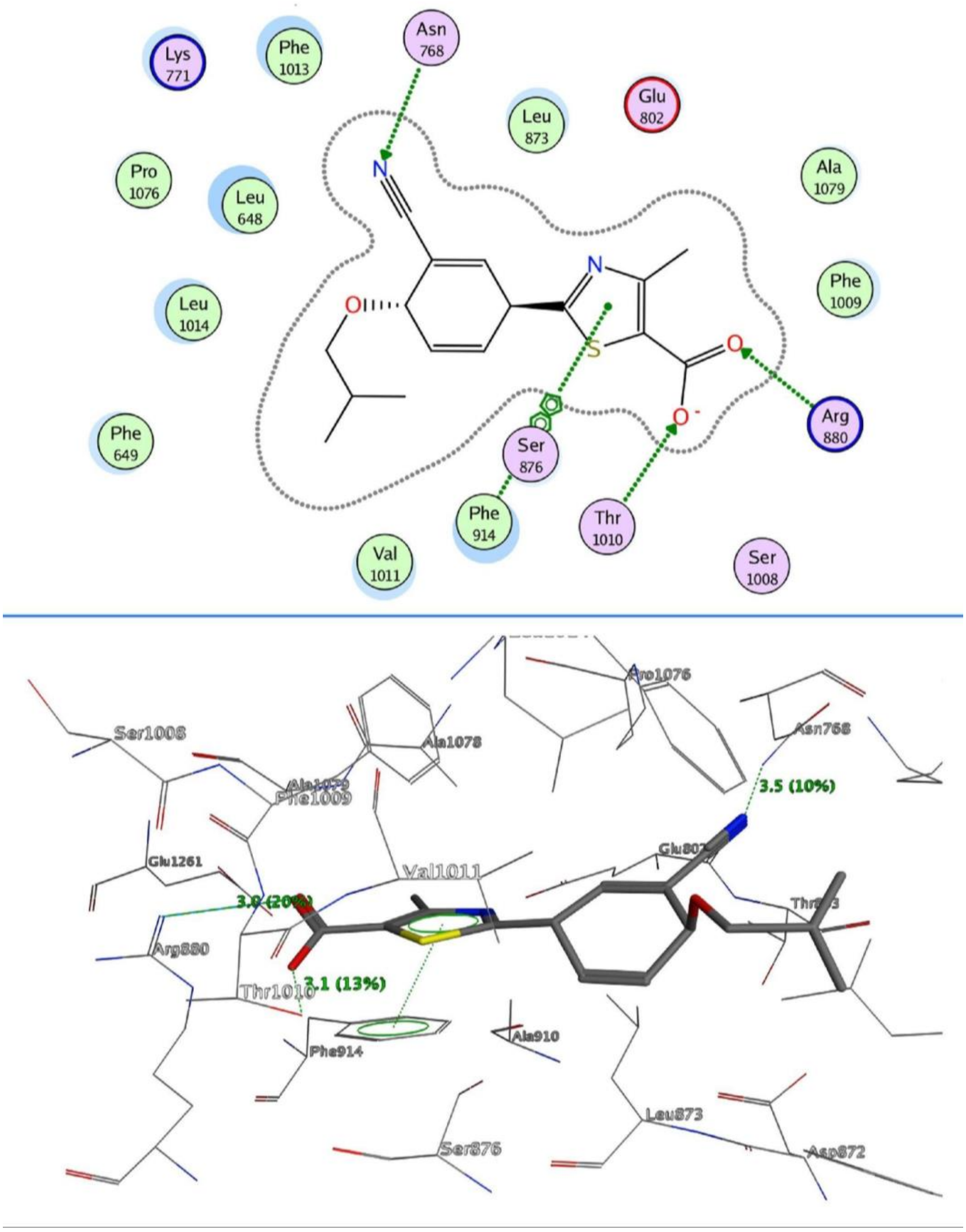


**Figure 5S.** 2D (upper) and 3D (lower) interactions of Xanthine Oxidase (PDB ID: 1N5X) with febuxostat (Fitness score of 64.53).

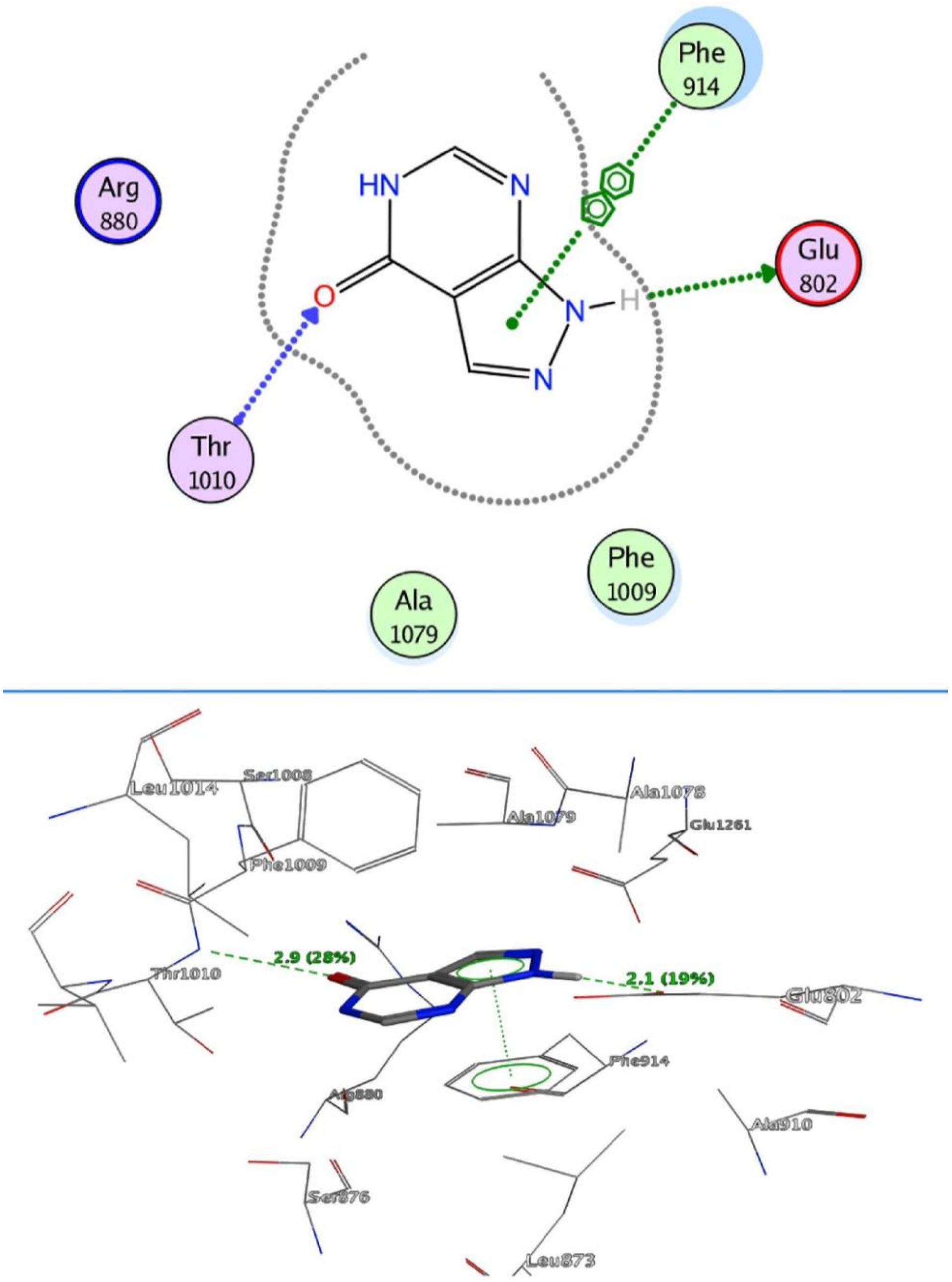


**Figure 6S.** 2D (upper) and 3D (lower) interactions of Xanthine Oxidase with allopurinol (Fitness score of 46.16).

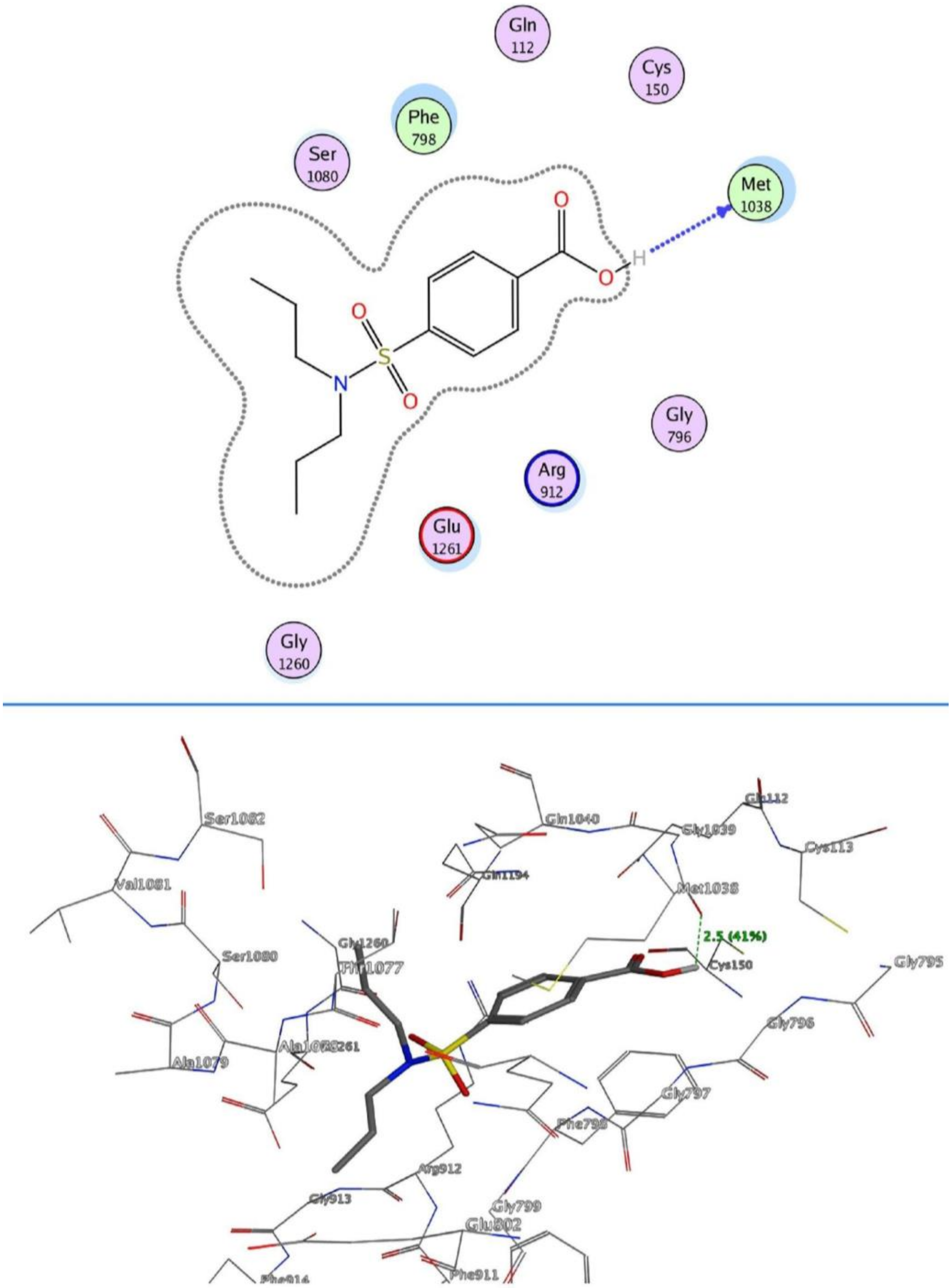


**Figure 7S.** 2D (upper) and 3D (lower) interactions of Xanthine Oxidase with probenecid (Fitness score of 57.75).

**References**


[1] K. Okamoto, B. T. Eger, T. Nishino, S. Kondo, E. F. Pai, and T. Nishino, "An Extremely Potent Inhibitor of Xanthine Oxidoreductase: CRYSTAL STRUCTURE OF THE ENZYME-INHIBITOR COMPLEX AND MECHANISM OF INHIBITION," *J. Biol. Chem.*, vol. 278, no. 3, pp. 1848–1855, Jan. 2003, doi: 10.1074/jbc.M208307200.

[2] E. S. Istvan, "Structural Mechanism for Statin Inhibition of HMG-CoA Reductase," *Science*, vol. 292, no. 5519, pp. 1160–1164, May 2001, doi: 10.1126/science.1059344.

[3] D. Iqbal, M. S. Khan, M. S. Khan, S. Ahmad, M. S. Hussain, and M. Ali, "Bioactivity guided fractionation and hypolipidemic property of a novel HMG-CoA reductase inhibitor from Ficus virens Ait," *Lipids Health Dis.*, vol. 14, no. 1, p. 15, Dec. 2015, doi: 10.1186/s12944-015-0013-6.

[4] J. Zhang *et al.*, "Eight new triterpenoids with inhibitory activity against HMG-CoA reductase from the medical mushroom Ganoderma leucocontextum collected in Tibetan plateau," *Fitoterapia*, vol. 130, pp. 79–88, Oct. 2018, doi: 10.1016/j.fitote.2018.08.009.

[5] K. Wang *et al.*, "Lanostane Triterpenes from the Tibetan Medicinal Mushroom *Ganoderma leucocontextum* and Their Inhibitory Effects on HMG-CoA Reductase and $\alpha$-Glucosidase," *J. Nat. Prod.*, vol. 78, no. 8, pp. 1977–1989, Aug. 2015, doi: 10.1021/acs.jnatprod.5b00331.

[6] K. Wang *et al.*, "A novel class of $\alpha$-glucosidase and HMG-CoA reductase inhibitors from Ganoderma leucocontextum and the anti-diabetic properties of ganomycin I in KK-A y mice," *Eur. J. Med. Chem.*, vol. 127, pp. 1035–1046, Feb. 2017, doi: 10.1016/j.ejmech.2016.11.015.

[7] B. Chen *et al.*, "Triterpenes and meroterpenes from Ganoderma lucidum with inhibitory activity against HMGs reductase, aldose reductase and $\alpha$-glucosidase," *Fitoterapia*, vol. 120, pp. 6–16, Jul. 2017, doi: 10.1016/j.fitote.2017.05.005.

[8] C. Li, Y. Li, and H. H. Sun, "New ganoderic acids, bioactive triterpenoid metabolites from the mushroom ***Ganoderma lucidum*** ," *Nat. Prod. Res.*, vol. 20, no. 11, pp. 985–991, Sep. 2006, doi: 10.1080/14786410600921466.

[9] M. Koo *et al.*, "3-Hydroxy-3-methylglutaryl-CoA (HMG-CoA) reductase inhibitory effect of Vitis vinifera," *Fitoterapia*, vol. 79, no. 3, pp. 204–206, Apr. 2008, doi: 10.1016/j.fitote.2007.11.005.

[10] E.-K. Kwon *et al.*, "Flavonoids from the Buds of *Rosa damascena* Inhibit the Activity of 3-Hydroxy-3-methylglutaryl-coenzyme A Reductase and Angiotensin I-Converting Enzyme," *J. Agric. Food Chem.*, vol. 58, no. 2, pp. 882–886, Jan. 2010, doi: 10.1021/jf903515f.

[11] F. Xu, X. Zhao, L. Yang, X. Wang, and J. Zhao, "A New Cycloartane-Type Triterpenoid Saponin Xanthine Oxidase Inhibitor from Homonoia riparia Lour," *Molecules*, vol. 19, no. 9, pp. 13422–13431, Aug. 2014, doi: 10.3390/molecules190913422.

[12] R. Arimboor, M. Rangan, S. G. Aravind, and C. Arumughan, "Tetrahydroamentoflavone (THA) from Semecarpus anacardium as a potent inhibitor of xanthine oxidase," *J. Ethnopharmacol.*, vol. 133, no. 3, pp. 1117–1120, Feb. 2011, doi: 10.1016/j.jep.2010.10.027.

[13] M. T. Nguyen, S. Awale, Y. Tezuka, J. Ueda, Q. L. Tran, and S. Kadota, "Xanthine Oxidase Inhibitors from the Flowers of *Chrysanthemum sinense*," *Planta Med.*, vol. 72, no. 1, pp. 46–51, Nov. 2006, doi: 10.1055/s-2005-873181.

[14] H.-X. Liu *et al.*, "Xanthine oxidase inhibitors isolated from Piper nudibaccatum," *Phytochem. Lett.*, vol. 12, pp. 133–137, Jun. 2015, doi: 10.1016/j.phytol.2015.03.005.

[15] A. NAGAO, M. SEKI, and H. KOBAYASHI, "Inhibition of Xanthine Oxidase by Flavonoids," *Biosci. Biotechnol. Biochem.*, vol. 63, no. 10, pp. 1787–1790, Jan. 1999, doi:

10.1271/bbb.63.1787.

[16] L.-N. Huo *et al.*, "Bioassay-Guided Isolation and Identification of Xanthine Oxidase Inhibitory Constituents from the Leaves of Perilla frutescens," *Molecules*, vol. 20, no. 10, pp. 17848–17859, Sep. 2015, doi: 10.3390/molecules201017848.

[17] S. Honda and T. Masuda, "Identification of Pyrogallol in the Ethyl Acetate-Soluble Part of Coffee as the Main Contributor to Its Xanthine Oxidase Inhibitory Activity," *J. Agric. Food Chem.*, vol. 64, no. 41, pp. 7743–7749, Oct. 2016, doi: 10.1021/acs.jafc.6b03339.

[18] D. Liu, D. Wang, W. Yang, and D. Meng, "Potential anti-gout constituents as xanthine oxidase inhibitor from the fruits of Stauntonia brachyanthera," *Bioorg. Med. Chem.*, vol. 25, no. 13, pp. 3562–3566, Jul. 2017, doi: 10.1016/j.bmc.2017.05.010.

[19] F. Nessa, Z. Ismail, and N. Mohamed, "Xanthine oxidase inhibitory activities of extracts and flavonoids of the leaves of *Blumea balsamifera*," *Pharm. Biol.*, vol. 48, no. 12, pp. 1405–1412, Dec. 2010, doi: 10.3109/13880209.2010.487281.

[20] T. Unno, A. Sugimoto, and T. Kakuda, "Xanthine oxidase inhibitors from the leaves of Lagerstroemia speciosa (L.) Pers.," *J. Ethnopharmacol.*, vol. 93, no. 2–3, pp. 391–395, Aug. 2004, doi: 10.1016/j.jep.2004.04.012.

[21] H. Yuk, Y.-S. Lee, H. Ryu, S.-H. Kim, and D.-S. Kim, "Effects of Toona sinensis Leaf Extract and Its Chemical Constituents on Xanthine Oxidase Activity and Serum Uric Acid Levels in Potassium Oxonate-Induced Hyperuricemic Rats," *Molecules*, vol. 23, no. 12, p. 3254, Dec. 2018, doi: 10.3390/molecules23123254.

[22] N. Masuoka, T. Isobe, and I. Kubo, "Antioxidants fromRabdosia japonica," *Phytother. Res.*, vol. 20, no. 3, pp. 206–213, Mar. 2006, doi: 10.1002/ptr.1835.

[23] B. S. Tuzun *et al.*, "Isolation of Chemical Constituents of Centaurea virgata Lam. and Xanthine Oxidase Inhibitory Activity of the Plant Extract and Compounds," *Med. Chem.*, vol. 13, no. 5, Jul. 2017, doi: 10.2174/1573406413666161219161946.

[24] C.-N. Lin *et al.*, "Xanthine oxidase inhibitory terpenoids of Amentotaxus formosana protect cisplatin-induced cell death by reducing reactive oxygen species (ROS) in normal human urothelial and bladder cancer cells," *Phytochemistry*, vol. 71, no. 17–18, pp. 2140–2146, Dec. 2010, doi: 10.1016/j.phytochem.2010.08.012.

[25] S. H. Nile and S. W. Park, "Antioxidant, $\alpha$-Glucosidase and Xanthine Oxidase Inhibitory Activity of Bioactive Compounds From Maize ( *Zea mays* L.)," *Chem. Biol. Drug Des.*, vol. 83, no. 1, pp. 119–125, Jan. 2014, doi: 10.1111/cbdd.12205.

[26] K.-W. Lin, S.-C. Yang, and C.-N. Lin, "Antioxidant constituents from the stems and fruits of Momordica charantia," *Food Chem.*, vol. 127, no. 2, pp. 609–614, Jul. 2011, doi: 10.1016/j.foodchem.2011.01.051.

[27] Y. Liu, S. Liu, and Z. Liu, "Screening and determination of potential xanthine oxidase inhibitors from Radix Salviae Miltiorrhizae using ultrafiltration liquid chromatography–mass spectrometry," *J. Chromatogr. B*, vol. 923–924, pp. 48–53, Apr. 2013, doi: 10.1016/j.jchromb.2013.02.009.

[28] S. H. Nile and S. W. Park, "Chromatographic analysis, antioxidant, anti-inflammatory, and xanthine oxidase inhibitory activities of ginger extracts and its reference compounds," *Ind. Crops Prod.*, vol. 70, pp. 238–244, Aug. 2015, doi: 10.1016/j.indcrop.2015.03.033.

[29] H.-Z. Jiang, S. Hu, R.-X. Tan, R. Tan, and R.-H. Jiao, "Neocucurbitacin D, a novel lactone-type norcucurbitacin as xanthine oxidase inhibitor from *Herpetospermum pedunculosum*," *Nat. Prod. Res.*, vol. 34, no. 12, pp. 1728–1734, Jun. 2020, doi: 10.1080/14786419.2018.1528592.

[30] C. X. Zhou, L. D. Kong, W. C. Ye, C. H. K. Cheng, and R. X. Tan, "Inhibition of Xanthine and Monoamine Oxidases by Stilbenoids from Veratrum taliense," *Planta Med.*, vol. 67, no.

2, pp. 158–161, 2001, doi: 10.1055/s-2001-11500.

[31] X. Tang, A. Xiao, S. Mei, P. Tang, L. Ren, and L. Liu, "Pueraria lobata Root Constituents as Xanthine Oxidase Inhibitors and Protective Agents against Oxidative Stress Induced in GES-1 Cells," *J. Braz. Chem. Soc.*, 2020, doi: 10.21577/0103-5053.20200108.

[32] N. S. Ahmad, M. Farman, M. H. Najmi, K. B. Mian, and A. Hasan, "Pharmacological basis for use of Pistacia integerrima leaves in hyperuricemia and gout," *J. Ethnopharmacol.*, vol. 117, no. 3, pp. 478–482, May 2008, doi: 10.1016/j.jep.2008.02.031.

[33] P. Phuwapraisirisan, P. Sowanthip, D. H. Miles, and S. Tip-pyang, "Reactive radical scavenging and xanthine oxidase inhibition of proanthocyanidins fromCarallia brachiata," *Phytother. Res.*, vol. 20, no. 6, pp. 458–461, Jun. 2006, doi: 10.1002/ptr.1877.

[34] Y.-H. Chu, C.-J. Chen, S.-H. Wu, and J.-F. Hsieh, "Inhibition of Xanthine Oxidase by *Rhodiola crenulata* Extracts and Their Phytochemicals," *J. Agric. Food Chem.*, vol. 62, no. 17, pp. 3742–3749, Apr. 2014, doi: 10.1021/jf5004094.

[35] K. Liu *et al.*, "Chemical Evidence for Potent Xanthine Oxidase Inhibitory Activity of Ethyl Acetate Extract of Citrus aurantium L. Dried Immature Fruits," *Molecules*, vol. 21, no. 3, Art. no. 3, Mar. 2016, doi: 10.3390/molecules21030302.

[36] K. Murata *et al.*, "Hydroxychavicol: a potent xanthine oxidase inhibitor obtained from the leaves of betel, Piper betle," *J. Nat. Med.*, vol. 63, no. 3, pp. 355–359, Jul. 2009, doi: 10.1007/s11418-009-0331-y.

[37] C. Xiao *et al.*, "Three new C-geranylated flavonoids from Paulownia catalpifolia T. Gong ex D.Y. Hong seeds with their inhibitory effects on xanthine oxidase," *Phytochem. Lett.*, vol. 36, pp. 162–165, Apr. 2020, doi: 10.1016/j.phytol.2020.02.002.

[38] R. H. Jiao, H. M. Ge, D. H. Shi, and R. X. Tan, "An Apigenin-Derived Xanthine Oxidase Inhibitor from *Palhinhaea c ernua*," *J. Nat. Prod.*, vol. 69, no. 7, pp. 1089–1091, Jul. 2006, doi: 10.1021/np060038a.

[39] P. Valentão, E. Fernandes, F. Carvalho, P. B. Andrade, R. M. Seabra, and M. L. Bastos, "Antioxidant Activity of *Centaurium erythraea* Infusion Evidenced by Its Superoxide Radical Scavenging and Xanthine Oxidase Inhibitory Activity," *J. Agric. Food Chem.*, vol. 49, no. 7, pp. 3476–3479, Jul. 2001, doi: 10.1021/jf001145s.

[40] K.-W. Lin, C.-H. Liu, H.-Y. Tu, H.-H. Ko, and B.-L. Wei, "Antioxidant prenylflavonoids from Artocarpus communis and Artocarpus elasticus," *Food Chem.*, 2009, Accessed: Jul. 12, 2020. [Online]. Available: https://agris.fao.org/agris-search/search.do?recordID=US201301605341.

[41] Z. Yu, W. P. Fong, and C. H. K. Cheng, *The dual actions of morin (3,5,7,2 ' ,4 ' -pentahydroxyflavone) as a hypouricemic agent: uricosuric effect and xanthine oxidase inhibitory activity*. 2006.

[42] L. D. Kong, Y. Zhang, X. Pan, R. X. Tan, and C. H. K. Cheng*, "Inhibition of xanthine oxidase by liquiritigenin and isoliquiritigenin isolated from Sinofranchetia chinensis:," *Cell. Mol. Life Sci.*, vol. 57, no. 3, pp. 500–505, Mar. 2000, doi: 10.1007/PL00000710.

[43] K.-W. Lin *et al.*, "Cytotoxic and antioxidant constituents from Garcinia subelliptica," *Food Chem.*, vol. 135, no. 2, pp. 851–859, Nov. 2012, doi: 10.1016/j.foodchem.2012.04.133.

[44] T. Wang, D. Li, B. Yu, and J. Qi, "Screening inhibitors of xanthine oxidase from natural products using enzyme immobilized magnetic beads by high-performance liquid chromatography coupled with tandem mass spectrometry," *J. Sep. Sci.*, vol. 40, no. 9, pp. 1877–1886, May 2017, doi: 10.1002/jssc.201601438.

[45] S. Y. Wang, C. W. Yang, J. W. Liao, W. W. Zhen, F. H. Chu, and S. T. Chang, "Essential oil from leaves of Cinnamomum osmophloeum acts as a xanthine oxidase inhibitor and

reduces the serum uric acid levels in oxonate-induced mice," *Phytomedicine*, vol. 15, no. 11, pp. 940–945, Nov. 2008, doi: 10.1016/j.phymed.2008.06.002.

[46] A. M. Ferrari, M. Sgobba, M. C. Gamberini, and G. Rastelli, "Relationship between quantum-chemical descriptors of proton dissociation and experimental acidity constants of various hydroxylated coumarins. Identification of the biologically active species for xanthine oxidase inhibition," *Eur. J. Med. Chem.*, vol. 42, no. 7, pp. 1028–1031, Jul. 2007, doi: 10.1016/j.ejmech.2006.12.023.

[47] P. Ayyappan and S. V. Nampoothiri, "Bioactive natural products as potent inhibitors of xanthine oxidase," in *Studies in Natural Products Chemistry*, vol. 64, Elsevier, 2020, pp. 391–416.

[48] L. Hartanti, S. M. K. Yonas, J. J. Mustamu, S. Wijaya, H. K. Setiawan, and L. Soegianto, "Influence of extraction methods of bay leaves (Syzygium polyanthum) on antioxidant and HMG-CoA Reductase inhibitory activity," *Heliyon*, vol. 5, no. 4, p. e01485, Apr. 2019, doi: 10.1016/j.heliyon.2019.e01485.

[49] A. O. Ademosun *et al.*, "Phenolics from grapefruit peels inhibit HMG-CoA reductase and angiotensin-I converting enzyme and show antioxidative properties in endothelial EA.Hy 926 cells," *Food Sci. Hum. Wellness*, vol. 4, no. 2, pp. 80–85, Jun. 2015, doi: 10.1016/j.fshw.2015.05.002.

[50] G. Baskaran, S. Salvamani, S. A. Ahmad, N. A. Shaharuddin, P. D. Pattiram, and M. Y. Shukor, "HMG-CoA reductase inhibitory activity and phytocomponent investigation of Basella alba leaf extract as a treatment for hypercholesterolemia," *Drug Des. Devel. Ther.*, vol. 9, pp. 509–517, Jan. 2015, doi: 10.2147/DDDT.S75056.

[51] S. Salvamani, B. Gunasekaran, M. Y. Shukor, N. A. Shaharuddin, M. K. Sabullah, and S. A. Ahmad, "Anti-HMG-CoA Reductase, Antioxidant, and Anti-Inflammatory Activities of Amaranthus viridis Leaf Extract as a Potential Treatment for Hypercholesterolemia," *Evidence-Based Complementary and Alternative Medicine*, 2016. https://www.hindawi.com/journals/ecam/2016/8090841/ (accessed Apr. 28, 2020).

[52] J. P.-C. Wong *et al.*, "Crude Ethanol Extract of *Pithecellobium ellipticum* as a Potential Lipid-Lowering Treatment for Hypercholesterolaemia," *Evid. Based Complement. Alternat. Med.*, vol. 2014, pp. 1–9, 2014, doi: 10.1155/2014/492703.

[53] K. A. Hafidz, N. Puspitasari, A. A, A. Yanuar, Y. Artha, and A. Mun'im, "HMG-CoA Reductase Inhibitory Activity of Gnetum gnemon Seed Extract and Identification of Potential Inhibitors for Lowering Cholesterol Level," *J. Young Pharm.*, vol. 9, no. 4, pp. 559–565, Oct. 2017, doi: 10.5530/jyp.2017.9.107.

[54] A. Gholamhose, B. Shahouzehi, and F. Sharifi-Fa, "Inhibitory Activity of Some Plant Methanol Extracts on 3-Hydroxy-3-Methylglutaryl Coenzyme a Reductase," *Int. J. Pharmacol.*, vol. 6, no. 5, pp. 705–711, May 2010, doi: 10.3923/ijp.2010.705.711.

[55] D. Iqbal *et al.*, "*In Vitro* Screening for $\beta$ -Hydroxy- $\beta$ -methylglutaryl-CoA Reductase Inhibitory and Antioxidant Activity of Sequentially Extracted Fractions of *Ficus palmata* Forsk," *BioMed Res. Int.*, vol. 2014, pp. 1–10, 2014, doi: 10.1155/2014/762620.

[56] Sulistiyani *et al.*, "Nrf2-inducing and HMG-CoA reductase inhibitory activities of a polyphenol-rich fraction of Guazuma ulmifolia leaves," *Asian Pac. J. Trop. Biomed.*, vol. 9, no. 9, p. 389, Sep. 2019, doi: 10.4103/2221-1691.267659.

[57] S. Rahmania and S. Sulistiyani, "Identification of HMG-CoA Reductase Inhibitor Active Compound in Medicinal Forest Plants," *J. Kefarmasian Indones.*, vol. 7, Aug. 2017, doi: 10.22435/jki.v7i2.6279.95-104.

[58] M. Umamaheswari, K. AsokKumar, A. Somasundaram, T. Sivashanmugam, V.

Subhadradevi, and T. K. Ravi, "Xanthine oxidase inhibitory activity of some Indian medical plants," *J. Ethnopharmacol.*, vol. 109, no. 3, pp. 547–551, Feb. 2007, doi: 10.1016/j.jep.2006.08.020.

[59] E. Abu-Gharbieh, N. G. Shehab, I. M. Almasri, and Y. Bustanji, "Antihyperuricemic and xanthine oxidase inhibitory activities of Tribulus arabicus and its isolated compound, ursolic acid: In vitro and in vivo investigation and docking simulations," *PLOS ONE*, vol. 13, no. 8, p. e0202572, Aug. 2018, doi: 10.1371/journal.pone.0202572.

[60] M. T. T. Nguyen, S. Awale, Y. Tezuka, Q. L. Tran, H. Watanabe, and S. Kadota, "Xanthine Oxidase Inhibitory Activity of Vietnamese Medicinal Plants," *Biol. Pharm. Bull.*, vol. 27, no. 9, pp. 1414–1421, 2004, doi: 10.1248/bpb.27.1414.

[61] M. Berboucha, K. Ayouni, D. Atmani, D. Atmani, and M. Benboubetra, "Kinetic Study on the Inhibition of Xanthine Oxidase by Extracts from Two Selected Algerian Plants Traditionally Used for the Treatment of Inflammatory Diseases," *J. Med. Food*, vol. 13, no. 4, pp. 896–904, Aug. 2010, doi: 10.1089/jmf.2009.0164.

[62] M. Hudaib *et al.*, "Xanthine oxidase inhibitory activity of the methanolic extracts of selected Jordanian medicinal plants," *Pharmacogn. Mag.*, vol. 7, no. 28, p. 320, 2011, doi: 10.4103/0973-1296.90413.

[63] L. D. Kong, Y. Cai, W. W. Huang, C. H. K. Cheng, and R. X. Tan, "Inhibition of xanthine oxidase by some Chinese medicinal plants used to treat gout," *J. Ethnopharmacol.*, vol. 73, no. 1–2, pp. 199–207, Nov. 2000, doi: 10.1016/S0378-8741(00)00305-6.

[64] J. Havlik *et al.*, "Xanthine oxidase inhibitory properties of Czech medicinal plants," *J. Ethnopharmacol.*, vol. 132, no. 2, pp. 461–465, Nov. 2010, doi: 10.1016/j.jep.2010.08.044.

[65] F. Abdulhafiz *et al.*, "Xanthine Oxidase Inhibitory Activity, Chemical Composition, Antioxidant Properties and GC-MS Analysis of Keladi Candik (Alocasia longiloba Miq)," *Molecules*, vol. 25, no. 11, p. 2658, Jun. 2020, doi: 10.3390/molecules25112658.
